# Sequentially Self-Assembled Supramolecular Nanocomplexes Enable Systemic Cas9 RNP Delivery and *In Vivo* Tumor Genome Editing

**DOI:** 10.64898/2026.05.08.723716

**Authors:** Takumi Matsuo, Yuto Honda, Toshizumi Chino, Takahiro Nomoto, Yuriko Osakabe, Yutaka Miura, Nobuhiro Nishiyama

**Author notes:** Corresponding author (Y.H.); (N.N.).

## Abstract

*In vivo* genome editing with CRISPR–Cas9 ribonucleoproteins (RNPs) holds substantial therapeutic promise, yet rapid bloodstream clearance and the absence of delivery systems capable of systemic tumor targeting have hindered its clinical translation. Herein, a supramolecular ternary complex platform is reported in which Cas9/sgRNA RNPs are co-assembled with tannic acid (TA) and phenylboronic acid (PBA)-conjugated polymers through sequential self-assembly, producing ∼30 nm core–shell ternary complexes that protect RNPs from enzymatic degradation and dissociate selectively at endosomal pH. Upon intravenous administration in subcutaneous tumor-bearing mice, these ternary complexes exhibit prolonged blood circulation and preferential tumor accumulation, achieving 37.2% gene editing at tumor sites compared with only 1.5% for free RNPs. The platform successfully knocks out previously undruggable oncogenes including mutant KRAS and polo-like kinase 1 (PLK1), markedly suppressing tumor growth *in vivo*. By integrating sequential supramolecular self-assembly with stimuli-responsive cargo release, this strategy establishes a generalizable framework for systemically administered *in vivo* CRISPR therapeutics.

## 1. Introduction

Clustered regularly interspaced short palindromic repeat (CRISPR)-associated protein 9 (Cas9) nuclease systems have emerged as a revolutionary gene editing tool with vast potential in biomedicine ^1–4^ and agriculture ^5^. Among the available formats, the direct delivery of preformed Cas9/single-guide-RNA (sgRNA) ribonucleoprotein (RNP) complexes is uniquely attractive because it bypasses transcription and translation, enables rapid genome editing, and minimizes off-target activity compared with plasmid-DNA, mRNA, or AAV-based approaches ^6,7,8^. These properties have motivated therapeutic development against cancer ^9^, diabetes ^10^, and muscular dystrophy ^11^.

Despite this promise, the clinical translation of RNP-based therapeutics has been hindered by two intrinsic barriers. First, the ∼12 nm size and slightly anionic surface of RNP complexes severely limit cell-membrane translocation and nuclear import. Second, as bacteria-derived assemblies, RNPs are highly susceptible to enzymatic degradation by RNases and proteases in the bloodstream, abolishing editing activity within minutes of intravenous administration. Cationic lipid nanoparticles ^12–14^, gold nanoparticles ^15–17^, and hyaluronic-acid-based carriers ^18^ have therefore been engineered to encapsulate RNPs and improve cytosolic delivery. However, most of these systems lack precise control over intracellular release and exhibit insufficient blood retention for systemic targeting; consequently, the majority of *in vivo* studies have relied on local injection or hepatic accumulation^14^. Critically, few studies have established a quantitative correlation between pharmacokinetic profiles and on-site editing efficiency, an unmet need that has constrained the rational design of systemic RNP carriers and delayed clinical approval of *in vivo* CRISPR therapeutics.

To address these challenges, we previously reported a systemically applicable, protein-loaded ternary nanocomplex formed by sequential self-assembly of cargo proteins with tannic acid (TA, Fig. 1A) and phenylboronic acid (PBA)-conjugated polymers(Fig. 1B) in aqueous media^19^. TA is a biocompatible and biodegradable polyphenol bearing multiple galloyl groups that engage proteins, viral capsids, and peptides through hydrogen bonding and hydrophobic interactions^20,21^. The same galloyl groups also form reversible boronate esters with PBA side-chain residues at physiological pH (7.4) but readily dissociate under endosomal acidity (pH ∼5.5) ^22,23^. This TA-boronate chemistry yields core–shell ternary nanocomplexes that exhibit prolonged blood circulation ^24^, preferential tumor accumulation ^19,25^, and acid-triggered cargo release with high spatiotemporal selectivity. To date, however, the platform has been validated only with native cargo proteins; whether it can preserve the structural integrity and *in vivo* editing competence of the much larger and enzymatically fragile Cas9 RNP has remained an open question.

**Figure. 1.**
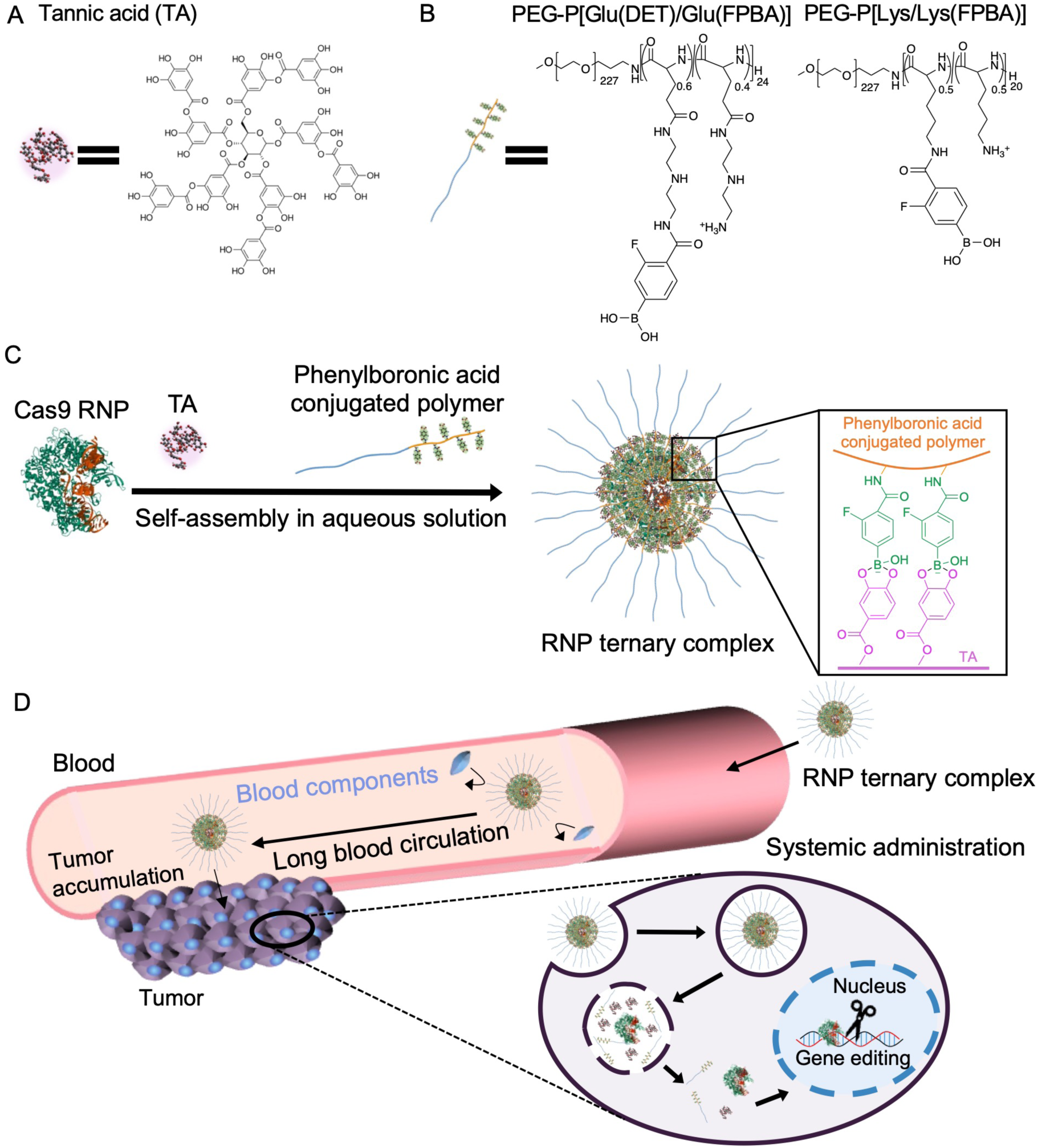
Concept of this study. Chemical structure of (A) TA and (B) PBA-conjugated polymer. (C) Schematic preparation of the ternary complex. The ternary complex is prepared through the sequential assembly of RNP, TA and PBA-conjugated polymers in aqueous solution. (D) Illustration of the RNP ternary complex and its potential function. The ternary complex prevents it from non-specific interaction with blood components, thus prolonged blood circulation and enhanced tumor accumulation were achieved without suppression of gene editing. After cellular uptake *via* endocytosis, the ternary complex releases the loaded RNP in response to endosomal acidic pH, thereby exhibiting efficient tumor gene editing.

In this study, we extend the supramolecular ternary complex platform to the systemic delivery of Cas9 RNPs and demonstrate, to our knowledge for the first time, a quantitative pharmacokinetics–editing relationship for an RNP carrier in a solid-tumor model (Figs. 1C,D). The resulting ∼30 nm RNP/TA/PBA-polymer ternary complexes exhibit prolonged blood circulation and preferential tumor accumulation following intravenous administration, achieve endosomal-pH-triggered RNP release, and reach 37.2% gene-editing efficiency at the tumor site versus 1.5% for free RNP. Leveraging this efficiency, we knock out previously undruggable oncogenes including mutant KRAS and polo-like kinase 1 (PLK1), markedly suppressing tumor growth *in vivo*. Together, these results establish a generalizable supramolecular framework for systemically administered *in vivo* CRISPR therapeutics that overcomes the tissue-targeting limitations of conventional RNP delivery systems.

## 2. Results

### 2.1. Preparation and characterization of RNP ternary complexes

Cas9/sgRNA RNP was prepared by mixing Cas9 protein with two nuclear localization sequences (2NLS) and sgRNA in aqueous solution. RNP was mixed with TA at various TA to RNP ratios ([TA]/[RNP]) in aqueous solution to prepare RNP/TA complexes. This [TA]/[RNP] ratio is an important indicator of complex formation because excess amounts of TA induce aggregation and precipitation of RNP owing to their hydrophobic nature ^21^. Based on a previous study ^19^, the formation of the RNP/TA complex was evaluated by light scattering analysis whereby the formation of a large aggregate was reflected as a significant increase in the scattered light intensity (derived count rate) (Fig. S1). For [TA]/[RNP] ≤ 30, the derived count rate barely increased with the addition of TA, whereas a steep increase in the derived count rate was observed at [TA]/[RNP] 2 50, suggesting distinct aggregation of RNP/TA complexes at [TA]/[RNP] ≤ 50. Thus, [TA]/[RNP] was set to 30 for all subsequent measurements.

We prepared two types of PBA-conjugated polymers: P[Glu(DET)/Glu(FPBA)] and P[Lys/Lys(FPBA)], in which 14 and 10 FPBA moieties were conjugated to side chains on the poly[Glu(DET)] component of PEG-P[Glu(DET)] block copolymer and on the poly(L-Lysine) component of PEG-PLys block copolymer, respectively, *via* amide linkage (M*_n_* of PEG:10,000, the DP of Glu(DET) and Lys: 24 and 20, respectively) (Figs. 1B, S2-S3 and Table S1). The FPBA moiety had a p*K*_a_ of 7.2, thus forming a stable boronate ester with the galloyl groups of TA at physiological pH (∼7.4) ^26^. In our previous study, P[Lys/Lys(FPBA)] was used to form protein ternary complexes with GFP/TA and b-gal/TA complexes, and the conjugation rate of FPBA to the Lys side chain was set to 50% to maintain the water solubility of the amine group residues of Lys, and the multiple FPBAs may permit the formation of multivalent boronate esters with the galloyl groups of TA ^19,25^. In addition, amine groups have been reported to improve the stability of nearby boronate esters through coordination bonding with boronic acid ^27^. Thus, to confirm the effect of the amine structures in the residues, P[Glu(DET)/Glu(FPBA)] with ethylenediamine moieties was used to prepare RNP/TA/PBA-conjugated polymer ternary complexes. The synthesis of each PBA-conjugated polymer was confirmed using GPC and ^1^H NMR spectroscopy (Figs. S4–S7).

P[Lys/Lys(FPBA)] or P[Glu(DET)/Glu(FPBA)] was mixed with the RNP/TA complex in aqueous solution, and the formation of RNP samples was evaluated by electrophoresis (Fig. 2A). To visualize the RNP samples, they were stained with Coomassie Brilliant Blue (CBB). Compared to RNP alone, the RNP/TA complexes exhibited longer migration toward the anode because of the anionic properties of TA, which is in good agreement with our previous studies ^19,25^. However, both RNP ternary complexes migrated to the cathode, suggesting that the RNP/TA complexes interact with PBA-conjugated polymers with cationic amine residues to form RNP ternary complexes. Interestingly, although the lanes of RNP/TA/P[Lys/Lys(FPBA)] contained only a spot, the lane of RNP/TA/P[Glu(DET)/Glu(FPBA)] contained a slight smear in addition to a spot, indicating a difference in the formation and stability of the RNP ternary complexes. Next, the hydrodynamic diameter and number of RNPs in each RNP complex were evaluated using fluorescence correlation spectroscopy (FCS) (Figs. 2B and C). Note that, this number of RNPs provides an average loading value, rather than indicating that every individual nanoparticle strictly contains one RNP molecule. The hydrodynamic diameter of RNP/TA was 48.5 nm, which is significantly larger than the RNP alone (13 nm) in aqueous solution, because a single RNP/TA complex contained ∼3 RNP particles, indicating that the RNP/TA complex was micro-aggregated due to the hydrophobic interaction of TA (Fig. 2C). After adding polymers to the RNP/TA complexes, the association number of the RNP ternary complexes decreased to ∼1.0, and the hydrodynamic diameters of the RNP ternary complexes with P[Glu(DET)/Glu(FPBA)] and P[Lys/Lys(FPBA)] changed to 21.7 nm and 27.9 nm, respectively, which are smaller than the RNP/TA complex but larger than single RNP. The DLS histogram revealed a single peak in all RNP samples (PDI of RNP, RNP/TA, and RNP/TA/P[Glu(DET)/Glu(FPBA)], and RNP/TA/P[Lys/Lys(FPBA)]: 0.35, 0.31, 0.29, and 0.13 respectively) (Fig. S8). Transmission electron microscopy (TEM) images showed that the RNP, RNP/TA/P[Glu(DET)/Glu(FPBA)], and RNP/TA/P[Lys/Lys(FPBA)] ternary complexes exhibited spherical and relatively dispersed morphologies, whereas RNP/TA showed micro aggregation in the dry state (Fig. S9). These results suggest that these polymers interacted with the RNP/TA complex and improved the dispersity owing to the hydrophilic properties of the polymers, strongly indicating the formation of RNP ternary complexes. Meanwhile, the hydrodynamic diameter of RNP/TA/P[Glu(DET)/Glu(FPBA)] was slightly smaller than that of RNP/TA/P[Lys/Lys(FPBA)], suggesting a difference in the stability and number of polymers in the complex, which is consistent with the electrophoresis results. To further investigate the differences between the polymers, the apparent affinity of the polymers for gallic acid (GA), which is a component molecule of TA, was measured by Alizarin red S (ARS) (Fig. 2D) ^22^. P[Lys/Lys(FPBA)] exhibited almost a 12-fold stronger apparent binding affinity for GA than P[Glu(DET)/Glu(FPBA)] because of the difference in the number of amines and the structure of the residual chain on the polymers. To summarize, both P[Glu(DET)/Glu(FPBA)] and P[Lys/Lys(FPBA)] could form RNP ternary complexes with the RNP/TA complex *via* boronate esters; however, P[Lys/Lys(FPBA)] interacted more strongly with the RNP/TA complex than P[Glu(DET)/Glu(FPBA)].

**Figure. 2.**
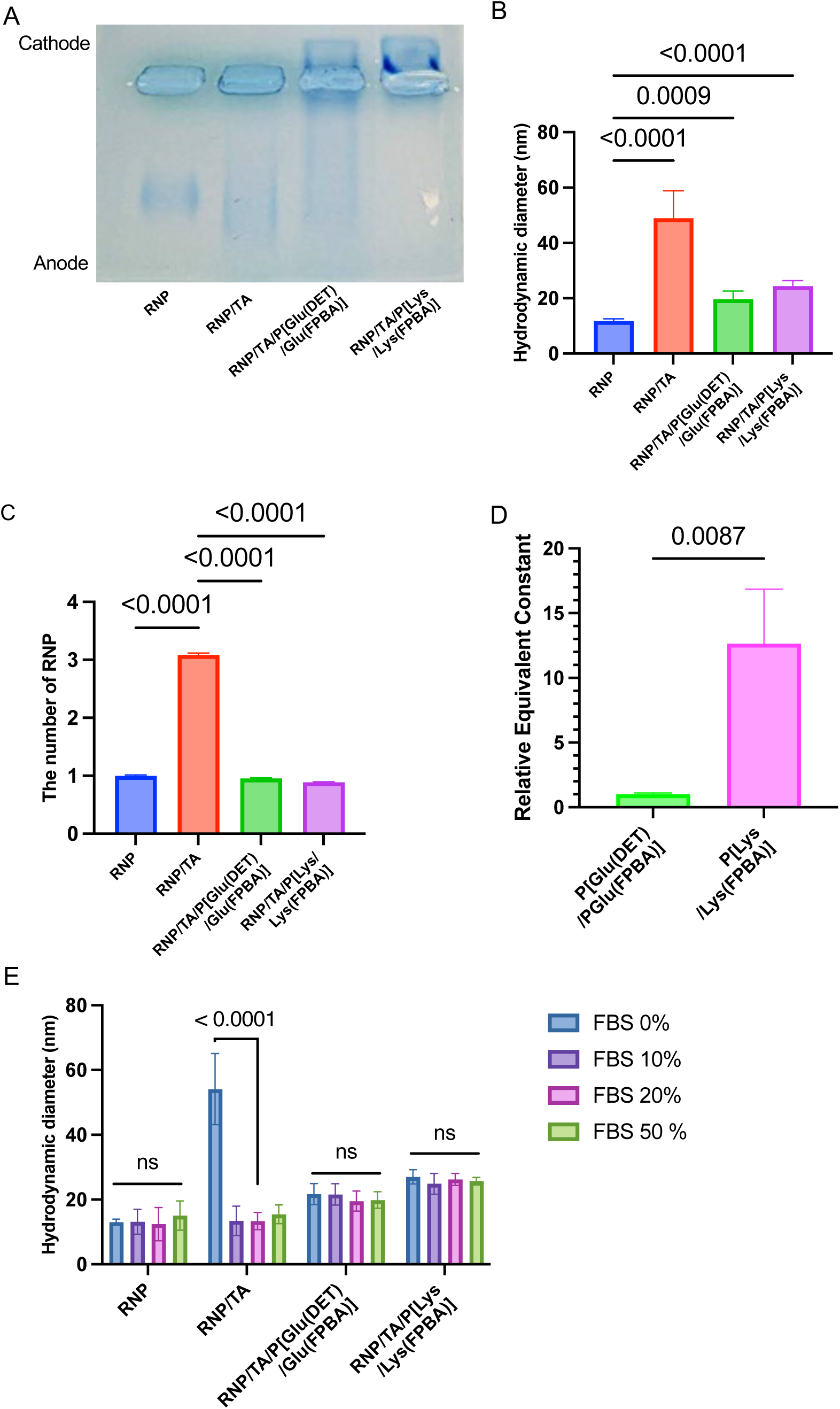
Characterization of RNP ternary complexes. (A) Native electrophoresis using agarose gel. These samples were stained with CBB. (B) Hydrodynamic diameters determined by FCS. The results are expressed as mean ± S.D. (n = 10). (C) The number of RNP in a complex determined by FCS. The results are expressed as mean ± S.D. (n = 10). (D) Relative apparent constant of the PBA-conjugated polymers with gallic acid. The relative apparent constant results are expressed as the relative values compared with the apparent constant of P[Glu(DET)/Glu(FPBA)]. The results are expressed as mean ± S.D. (n = 3). (E) Stability in FBS mixed solutions. Cas9 with GFP tag was used. The results are expressed as mean ± S.D. (n = 10). RNP concentration: 200 nM, [RNP]/[TA]/[PBA-conjugated polymer]= 1/30/30. ANOVA with Tukey’s multiple comparison test was used for statistical analysis in (B), (C), and (E). Unpaired t-test was used for statistical analysis in (D).

### 2.2. Stability of RNP ternary complexes under biological conditions

Since proteins in the blood, such as albumin, are known to interact with TA and protein/TA complexes ^28^, we used FCS to measure the hydrodynamic diameter of the RNP samples in the presence of FBS (Fig. 2E). The hydrodynamic diameter of the RNP was not significantly changed with increasing FBS concentrations, while the hydrodynamic diameter of the RNP/TA complex decreased from 49.0 nm to 12.2 nm when the FBS concentration was increased to 10%. It can be assumed that serum proteins in FBS interact with excess TA in the micro-aggregated RNP/TA complex and partially detach TA from the RNP/TA complex. Upon further increase of the FBS concentration to 50%, the diameter of the RNP/TA complex gradually increased to 14.0 nm, indicating that the serum components gradually interacted with RNP/TA complex. On the other hand, the hydrodynamic sizes of both RNP ternary complexes were stable even in the presence of 50% FBS. This behavior indicates that the PEG in PBA-conjugated polymers likely functioned as a shell and suppressed the interaction of the RNP/TA complex with FBS, consistent with our previous study ^19^. The stability of the RNP complexes was further confirmed by electrophoresis following incubation with FBS (Fig. S10). After incubation of each RNP sample with different concentration of FBS in aqueous solution at 37 °C for 24 hours, the RNP samples were electrophoresed, and the sgRNAs were stained with SYBR™ green II. In the presence of FBS, the RNP-derived band disappeared, and only the FBS-derived smear band was observed, suggesting that FBS components may degrade sgRNA. Meanwhile, both RNP ternary complexes exhibited bands of equivalent position and intensity to the untreated RNP ternary complex even in the presence of FBS. These results strongly suggest that the RNP ternary complex maintains stable particle formation and can protects the core RNP even in serum environments without apparent interference from the protein corona.

Next, we evaluated the resistance of the RNP samples to enzymatic degradation in the presence of RNase by electrophoresis (Fig. S11). RNase is present in the body and degrades sgRNAs of RNPs administered *in vivo*, resulting in decreased gene editing efficacy ^29^. After incubation of each RNP sample with RNase in aqueous solution, the RNP samples were electrophoresed, and the sgRNAs were stained with SYBR™ green II. The stained sgRNA bands of RNPs and RNP/TA complexes disappeared after RNase treatment, indicating degradation of the loaded sgRNA. In contrast, in both ternary complex lanes, the RNase-treated samples had band intensities similar to those of the untreated samples. This can be explained by the ternary complexes preventing RNase from accessing the core of the RNPs owing to the exclusion volume effect of the PEG shell. These results strongly suggest that P[Lys/Lys(FPBA)] and P[Glu(DET)/Glu(FPBA)] were sufficient to cover the RNP/TA complex.

The pH responsiveness of P[Glu(DET)/Glu(FPBA)] and P[Lys/Lys(FPBA)] was compared by measuring the hydrodynamic diameter of RNP ternary complexes in various pH solutions using FCS. The complexes exhibited a size reduction in response to decreasing pH, eventually reaching the size of RNP itself at pH 5.5 (Fig. S12). This behavior is consistent with previous studies conducted with GFP and β-gal ^19,25^.

### 2.3. Cellular uptake and subcellular distribution

Cellular uptake was quantified by measuring the fluorescence intensity of Cas9-GFP in HeLa-Luc cells by flow cytometry (FCM) (Fig. 3A). The cellular uptake of the RNP/TA complex was decreased to 40% of that of the RNPs, suggesting that serum components in the medium may detach TA from the RNP/TA complex, and the serum/TA complex might compete with the RNP/TA complex for interaction with the cell membrane, thereby inhibiting the cellular uptake of the RNP/TA complex. Both RNP/TA/P[Glu(DET)/Glu(FPBA)] and RNP/TA/P[Lys/Lys(FPBA)] ternary complexes also showed a decreased cellular uptake, but RNP/TA/P[Lys/Lys(FPBA)] ternary complexes demonstrated much lower cellular uptake, because RNP/TA/P[Lys/Lys(FPBA)] ternary complexes has more stable PEG-shell coating than RNP/TA/P[Glu(DET)/Glu(FPBA)] (Figs. 2A, and 2B), suppressing their interaction with cell membrane ^30^. Although flow cytometry suggested relatively high cellular association of naked RNPs, it should be noted that flow cytometry quantifies total cellular fluorescence and does not distinguish between membrane-bound and internalized materials. Therefore, biomolecules that strongly adsorb to the cell membrane can produce high fluorescence signals without efficient intracellular delivery. To clarify this point, the intracellular localization of each sample was further examined by confocal laser scanning microscopy.

**Figure. 3.**
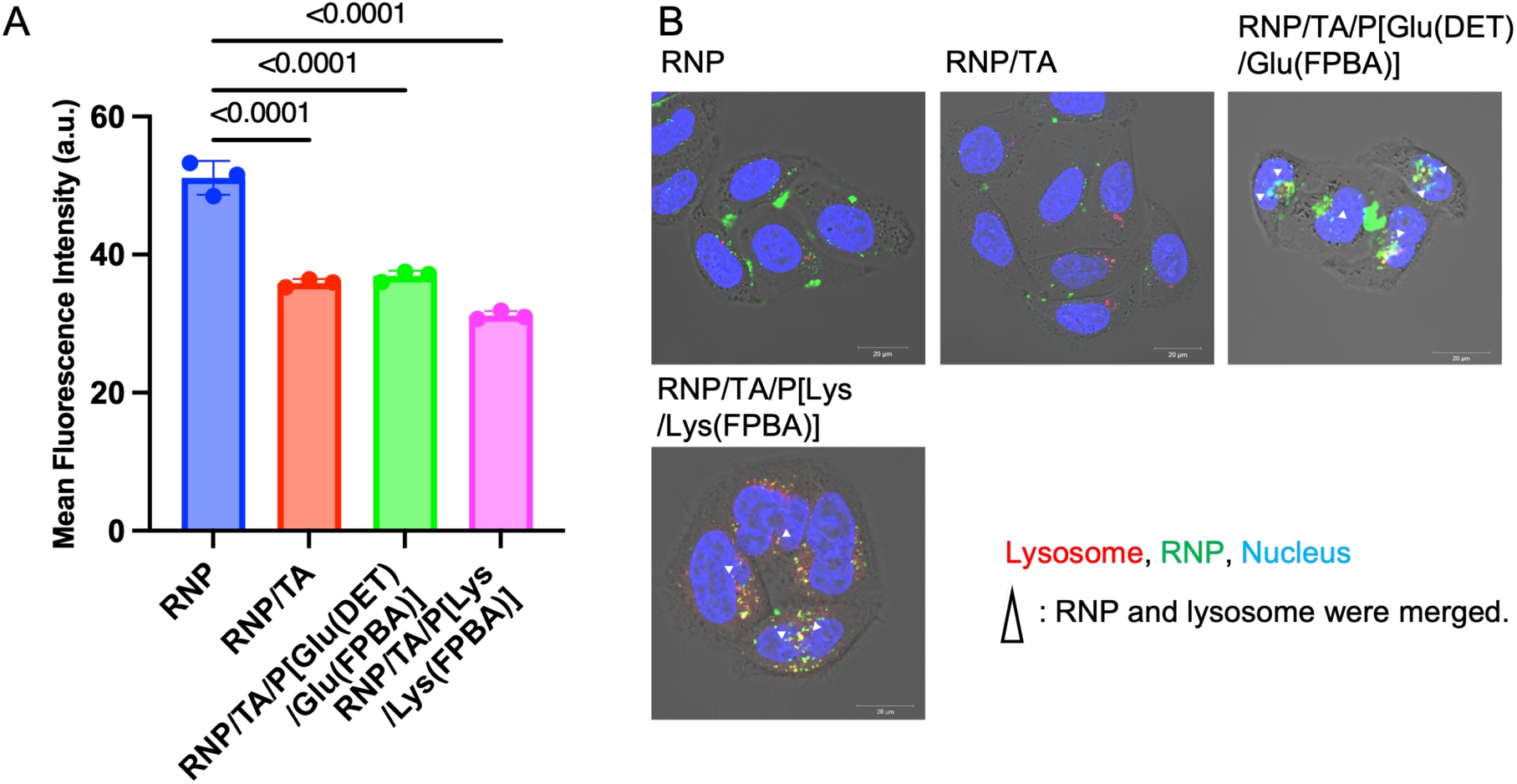
Subcellular distribution and cellular uptake of RNP samples. Cas9 with GFP tag was used. RNP concentrations: 60 nM, [RNP]/[TA]/[PBA-conjugated polymer]= 1/30/60. (A) Cellular uptake in HeLa-Luc cells evaluated by FCM. The cells were incubated with the samples for 24 h. The results are expressed as mean ± S.D. (n = 3). ANOVA with Tukey’s multiple comparison test was used for statistical analysis. (B) Subcellular distribution observed by CLSM. The samples were incubated with the samples for 24 h. RNP, Lysotracker Red DND-99, and nucleus are shown in green, red, and blue respectively. Scale bar, 10 μm.

The subcellular distribution of each RNP sample was observed in HeLa–Luc cells using Confocal Laser Scanning Microscopy (CLSM) 24 hours after sample treatment, and endo/lysosomes and nuclei were stained with LysoTracker™ Red DND-99 (red) and Hoechst 33342 (Blue), respectively (Fig. 3B, Fig. S13). CLSM observations revealed that the RNPs and RNP/TA complexes (green) were localized with the cell membrane, similar to a previous study ^31^. In contrast, the green fluorescence of both RNP ternary complexes not only overlapped with the red fluorescence (LysoTracker) and turn to yellow, but also diffused throughout the cells and overlapped with the blue (nucleus) fluorescence. This indicates that the polymer coating facilitated the endocytosis-mediated internalization of the RNP complexes, followed by their escape into the cytosol and subsequent translocation to the nucleus. Furthermore, the subcellular distribution images of RNP ternary complexes containing Cas9-GFP (green) and Alexa647-labelled PBA-conjugated polymers (orange) indicated that the intracellular distribution behavior of RNP and PBA-conjugated polymer did not correlate, and the Pearson correlation coefficient for both ternary complexes was approximately 0.1 (Fig. S14). Generally, the Pearson correlation coefficient below 0.2 indicates no association between the behavior of the two fluorophores, suggesting that RNP has dissociated from PBA-conjugated polymers within cells.

### 2.4. *In vitro* gene knockout study

To evaluate the gene knockout efficiency of the RNP ternary complexes, we incubated HeLa–Luc cells with RNP samples containing luciferase (Luc)-targeted sgRNA (Table. S2). The sgRNA sequence exhibited Luc gene editing in the previous study ^14^. After 72 hours of incubation, the luminescence intensity of treated cells was measured using a luminometer. (Fig. 4). According to the Cell counting kit-8 (CCK-8) assay, RNP/TA/P[Lys/Lys(FPBA)] ternary complexes showed slight toxicity compared to RNP/TA/P[Glu(DET)/Glu(FPBA)]. This might be because the primary amine structure in P[Lys/Lys(FPBA)] disrupted the cell membrane. To see the gene knockout efficiency, the luminescence intensity was normalized to the cell’s viability. The cells treated with RNP, or the RNP/TA complex exhibited similar luminescence intensity as D-PBS(-)-treated cells, indicating that RNP alone and the RNP/TA complex did not achieve adequate gene knockout in HeLa–Luc cells because of localization on the cell membrane and low nuclear transition. By contrast, RNP/TA/P[Glu(DET)/Glu(FPBA)] and RNP/TA/P[Lys/Lys(FPBA)] ternary complexes showed 13% and 40% decreases, respectively, in the luminescence intensities of the treated cells, indicating that the coating of the RNP/TA complexes with PBA-conjugated polymers augmented the gene knockout efficiency because of the enhanced intracellular uptake and nuclear translocation. The reduction of Luc expression was further confirmed by Western blot (Fig. S15). Compared to PBS-treated group, the signal was weaker in RNP/ProDeliverIN-treated cells and both RNP ternary complexes-treated cells, which is consistent with the result of Luc activity assay (Fig. 4A). The RNP/TA/P[Lys/Lys(FPBA)] ternary complex exhibited high gene knockout efficiency in Fig. 4A despite similar intracellular uptake of RNP/TA/P[Glu(DET)/Glu(FPBA)]. This may be because the RNP/TA/P[Lys/Lys(FPBA)] complex was covered by more polymer chains than the RNP/TA/P[Glu(DET)/Glu(FPBA)] complex owing to its high stability. Hence, a large number of P[Lys/Lys(FPBA)] chains may affect the endosomal membrane, resulting in high gene editing efficiency.

**Figure. 4.**
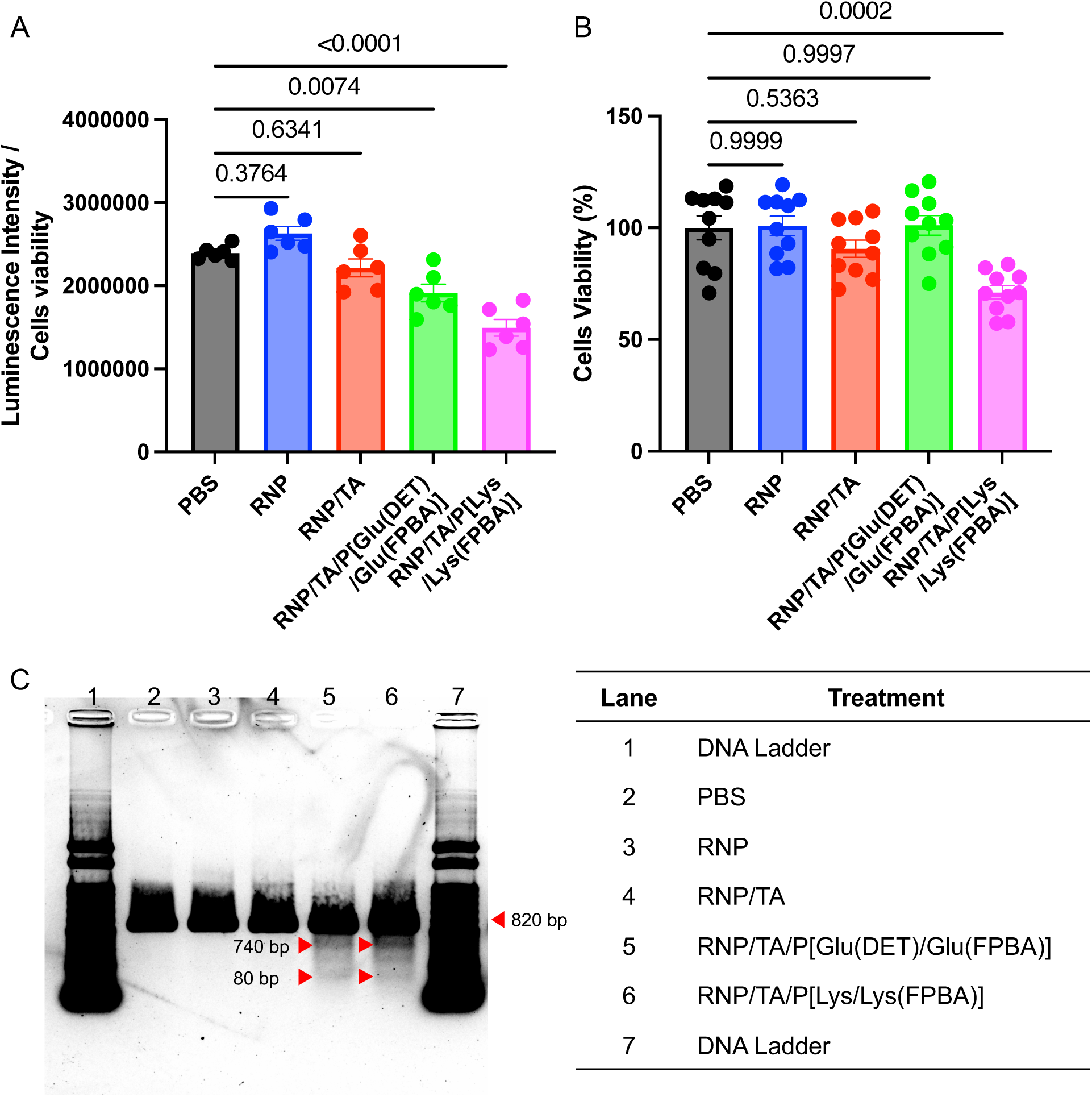
Luc gene editing by RNP ternary complex *in vitro*. RNP concentrations: 30 nM, [RNP]/[TA]/[PBA-conjugated polymer]= 1/30/60. (A) Luminescence intensity measurement. The cells were incubated with RNP samples for 72 h. The results are expressed as mean ± S.E.M. (n = 6) and normalized by cell’s viability. (B) Cytotoxicity assay. The cells were incubated with RNP samples for 72 h. The results are expressed as mean ± S.D. (n = 10). (C) T7EI cleavage assay of DNA isolated from HeLa-Luc cells treated with various RNP samples. Red arrows indicate cleavage bands. This experiment was repeated three times independently with similar results. ANOVA with Tukey’s multiple comparison test was used for statistical analysis in (A) and (B).

Knockout of Luc gene was further confirmed by T7 Endonuclease I (T7EI) assay (Fig. 4C). T7EI assay demonstrated that target DNA bands (820 bp) were cut into two cleavage bands (740 bp and 80 bp) at the ternary complex treated group, and no cleavage bands were observed with D-PBS(-)-treated group. In addition, to evaluate *in vitro* gene editing ability of RNP ternary complexes, we mixed RNP samples with an extracted and amplified Luc target DNA at physiological pH (∼7.4) or endosomal acidic pH (∼5.5), and the cleaved DNA was analyzed by gel electrophoresis. (Fig. S16). The DNA cleavage capacity of an RNP alone was equivalent at pH 5.5 and pH 7.4. Meanwhile, both RNP ternary complexes showed a single band derived from substrate DNA at pH 7.4, but showed 3 different bands derived from DNA cleaved into longer and shorter fragments at pH 5.5, indicating that RNP ternary complexes can release the loaded RNP in response to the endosomal acidic pH and recover the gene knockout function within the cells.

### 2.5. Biodistribution of the RNP ternary complexes

To examine the biodistribution profile of the RNP ternary complexes, we intravenously injected Cy5-labeled sgRNA alone and each RNP sample, (Cas9: 32.0 μg/mouse, sgRNA: 6.4 µg/mouse), into BALB/c nude mice bearing HeLa-Luc tumors, and the amounts of Cy5-labeled sgRNA in the tumors and normal organs were quantified using fluorophotometer (Fig. 5). The blood retention levels of RNP were 6.5% and 0.4% ID/mL plasma at 1 and 2 h after intravenous injection, respectively, whereas those of sgRNA alone were only 2.8% and 0.1% ID/mL plasma at 1 and 2 h, respectively (Fig. 5A). In addition, accumulation of the sgRNA alone in liver (0.8% ID/g liver) and kidney (1.3% ID/g kidney) at 2 h was lower than that of the RNP (6.0% ID/g liver and 6.0% ID/g kidney) (Figs. 5C and 5G). These differences indicate that stable RNP complexes formed even *in vivo*. The RNP/TA complexes had values of 2.5% and 0.7% ID/mL in plasma after 1 and 2 h, respectively, indicating a similar blood retention profile to RNP alone (Fig. 5A). However, the RNP alone exhibited considerable accumulation in liver and kidney (12.7% and 7.8% ID/g organ, respectively), whereas the RNP/TA complex exhibited high accumulation in liver and spleen (9.5% and 14.3% ID/g organ, respectively), with only 0.67% ID/g organ in kidney, likely owing to the TA-coating effect ^25^ (Figs. 5C and 5D). On the other hand, both RNP ternary complexes exhibited prolonged blood retention compared to the RNP alone and the RNP/TA complex. The blood retention values of the RNP/TA/P[Glu(DET)/Glu(FPBA)] ternary complexes were 29.9, 21.3, and 12.2% ID/mL plasma at 1, 2, and 6 h, respectively, and those of the RNP/TA/P[Lys/Lys(FPBA)] were 15.9, 10.1, and 5.6% ID/mL plasma at 1, 2, and 6 h, respectively (Fig. 5A). Such prolonged blood retention was probably due to the PEG shell, which prevented interaction with blood components, and the substantial coverage of RNP/TA by both polymers. Incidentally, that of the RNP/TA/P[Glu(DET)/Glu(FPBA)] ternary complex was approximately twice that of the RNP/TA/P[Lys/Lys(FPBA)] ternary complex. The organ accumulation profiles were also different, with the RNP/TA/P[Glu(DET)/Glu(FPBA)] ternary complex exhibiting high accumulation in kidney (46.6% ID/g kidney at 2 h), whereas the RNP/TA/P[Lys/Lys(FPBA)] ternary complex exhibited low accumulation in kidney (4.5% ID/g kidney at 2 h) (Figs. 5E and 5F). As an alternative to accumulation in kidney, the RNP/TA/P[Lys/Lys(FPBA)] ternary complex mainly accumulated in liver (36.6% ID/g liver at 2 h). This difference may be explained by differences in the structural stability of the ternary complexes (Figs. 5E and 5F). As shown in Fig. 2D, P[Lys/Lys(FPBA)] exhibited a stronger apparent affinity for GA than P[Glu(DET)/Glu(FPBA)], suggesting that the RNP/TA/P[Lys/Lys(FPBA)] ternary complex may be tightly covered by more polymer chains and exhibit more structural stability than the RNP/TA/P[Glu(DET)/Glu(FPBA)] ternary complex. The zeta-potential values of both TA/PBA-conjugated polymers were comparable, and the difference in the density of PBA-conjugated polymers in the ternary complexes may cause accumulation in the liver (Fig. S17). Note that, both RNP ternary complexes exhibited comparably low accumulation in spleen (≤ 10% ID/g spleen).

**Figure. 5.**
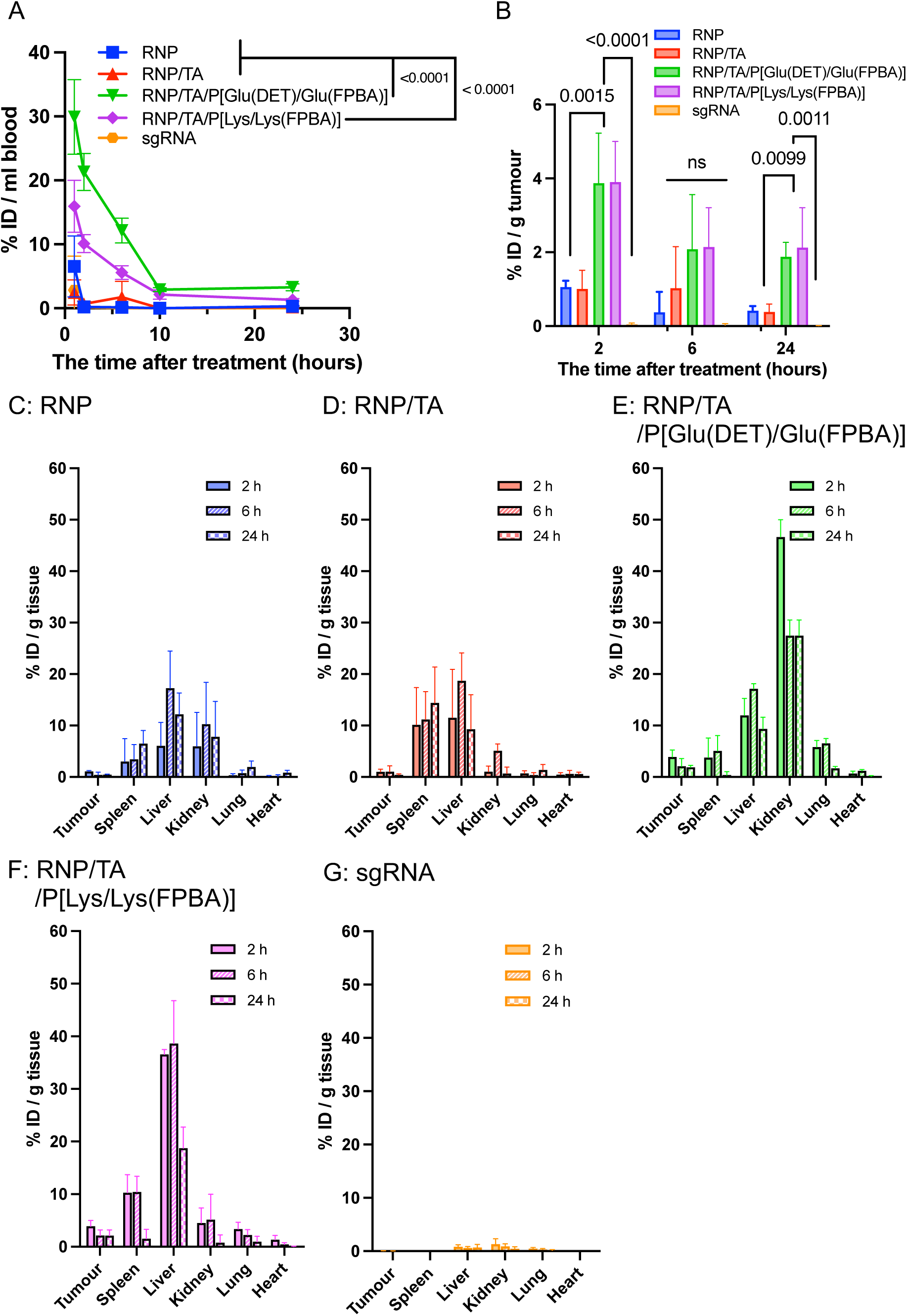
Biodistribution of intravenously injected RNP samples (32.0 μg of Cas9 and 6.4 μg) in subcutaneous HeLa-Luc tumor models. (A) Blood retention. The statistical difference at 2 hours post treatment was shown. (B) Tumor accumulation. (C-G) Biodistribution of (C) RNP, (D) RNP/TA, (E) RNP/TA/P[Glu(DET)/Glu(FPBA)], (F) RNP/TA/P[Lys/Lys(FPBA)] and (G) sgRNA. All the values were determined by Cy5 labelled sgRNA by fluorophotometer. The results are expressed as mean ± S.D. (n = 4). ANOVA with Tukey’s multiple comparison test was used for statistical analysis in (A) and (B).

Regarding tumor accumulation, the RNP and RNP/TA complex exhibited accumulation of approximately 1.0% ID/g tumor. On the other hand, both RNP ternary complexes exhibited 2%–4% ID/g tumor accumulation, achieving higher tumor accumulations than RNP and RNP/TA (Fig. 5B). These tumor accumulation properties are consistent with those of our previous studies ^19,25^. This significantly enhanced tumor accumulation may be due to an enhanced permeability and retention (EPR) effect of the ternary complex.

### 2.6. *In vivo* gene knockout study

To evaluate the *in vivo* gene editing efficiency, we intravenously injected each RNP sample (Cas9: 32.0 μg/mouse, sgRNA: 6.4 µg/mouse) containing Luc-targeted sgRNA into BALB/c nude mice bearing HeLa-Luc tumors. The luminescence intensities of the tumors were measured semi-quantitatively using an *in vivo* imaging system (IVIS) before sample injections and 3 and 7 days after sample injections (Figs. 6A and B). The luminescence intensity of each mouse on day 0 was set to 1 to calculate the time-dependent changes in the luminescence intensities (Fig. 6C). Note that the injection of RNP samples had no effect on tumor size growth (Fig. S18). The luminescence intensities at the tumor sites of RNP-and RNP/TA-treated mice were slightly lower than those of D-PBS(-)-treated mice at both 3 and 7 days, indicating the *in vivo* gene knockout efficiency of RNP. However, both RNP ternary complexes significantly suppressed the luminescence intensities of the tumors compared with other RNP samples on days 3 and 7, which can be attributed to high tumor accumulation. The luminescence intensities at the tumor sites of mice treated with the RNP ternary complexes on day 7 decreased compared to those on day 3, which suggests that the polymer coated RNP ternary complex was gradually translocated to the cell nucleus, possibly because of the modulated cellular uptake and transition to the cytoplasm.

**Figure. 6.**
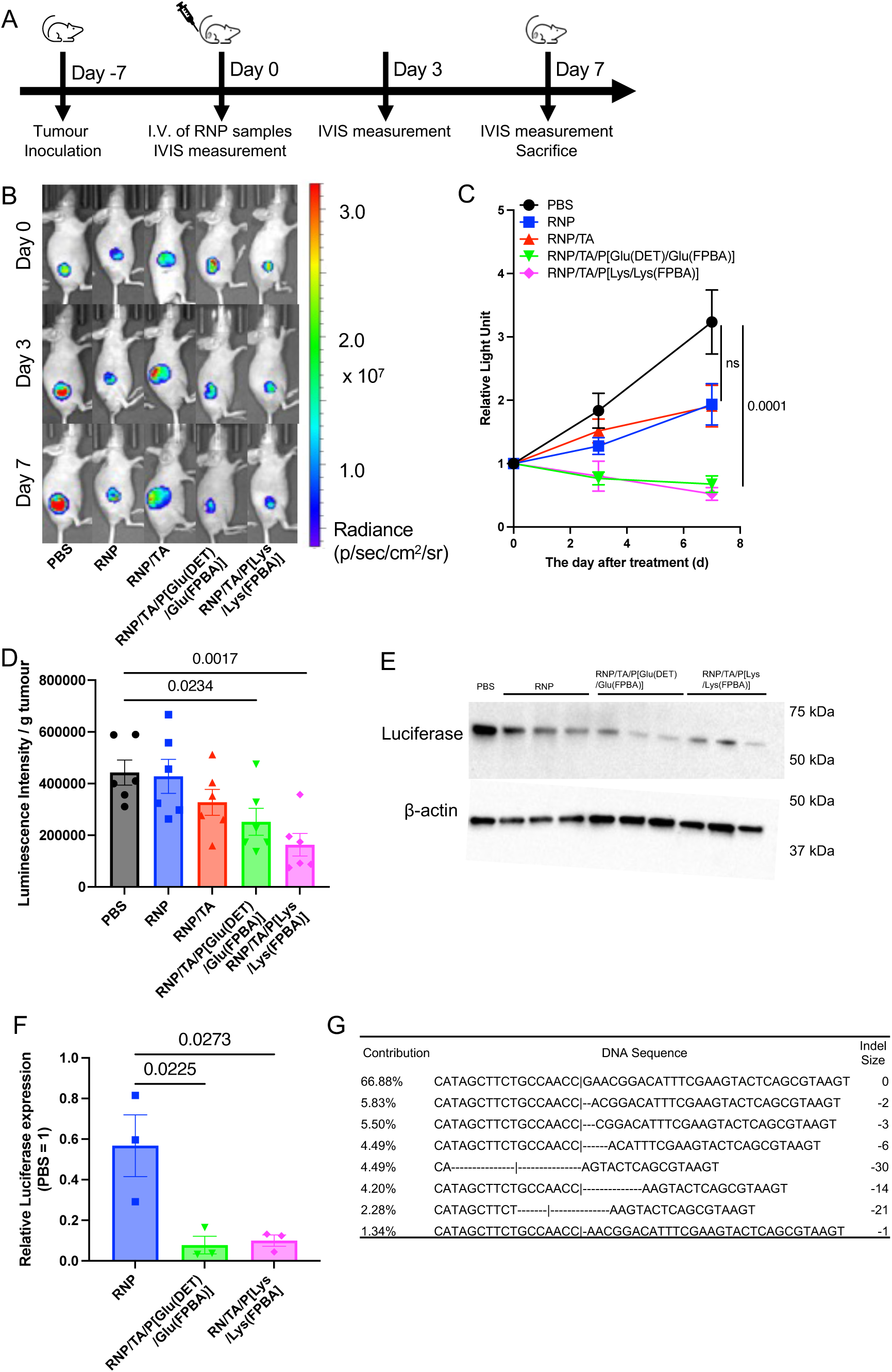
Luc gene editing by intravenously injected RNP samples (32.0 μg of Cas9 and 6.4 μg of sgRNA/mouse) in subcutaneous HeLa-Luc tumor models. (A) Scheme of the experiment. (B) The photos of mice received treatments taken by IVIS. (C) Luminescence intensity of the HeLa-Luc tumor. The value is normalized by the luminescence intensity on day 0. (D) Luminescence intensity of the lysed tumor. The intensity was normalized by the tumor weight. (E) Representative Western blot image showing Luc expression in Hela-Luc tumors treated with PBS, RNP, RNP/TA/P[Glu(DET)/Glu(FPBA)] and RNP/TA/P[Lys/Lys(FPBA)]. (F) Densitometric quantification of Luc levels normalized to Actin levels, and the relative values were then plotted against the PBS-treated group. (G) Sanger sequencing results of T-A cloning from HeLa-Luc tumor tissue after RNP/TA/P[Lys/Lys(FPBA)] treatment. The results are expressed as mean ± S.E.M. (n = 5 or 6). ANOVA with Tukey’s multiple comparison test was used in (C) and (F), and unpaired two-tailed t-tests with Welch correction was used in (D).

For a more quantitative evaluation, tumors were collected and homogenized on day 7, and the luminescence intensity of the tumors was measured by Luc assay (Fig. 6D). The RNP and RNP/TA exhibited a slight decrease in the luminescence intensities of the homogenized tumors to 96 % and 74 %, respectively, compared to the treatment with D-PBS(-), and the tendency was similar with the luminescence intensity from IVIS. Similar to the trend in luminescence intensity with IVIS, the RNP/TA/P[Glu(DET)/Glu(FPBA)] and RNP/TA/P[Lys/Lys(FPBA)] ternary complexes have exhibited clear gene editing effects and significantly decreased the luminescence intensities of the tumors to 54% and 36%, respectively, compared to that of D-PBS(-). In the luminescence measurement, the *in vivo* gene-editing effect of the RNP/TA/P[Lys/Lys(FPBA)] ternary complex was higher than that of the RNP/TA/P[Glu(DET)/Glu(FPBA)] ternary complex, although the tumor accumulation was comparable. The difference in gene editing effects can be explained by the higher *in vitro* gene knockout efficiency of RNP/TA/P[Lys/Lys(FPBA)] (Fig. 4A), which suggests that the RNP/TA/P[Lys/Lys(FPBA)] ternary complex is more effective as a systemically administered RNP delivery system against tumors. To analyze the Luc expression level in the tumors, RNP samples were administrated intravenously 3 times into HeLa-Luc tumor model mice. At 12 days post RNP samples injection, the tumors were analyzed by Western blot and were fixed with paraformaldehyde (PFA) for immunofluorescence staining. (Fig. 6E, 6F, S16). The Western blot image demonstrated that RNP treatment alone slightly reduced the Luc expression compared with non-treatment, while treatment with both RNP ternary complexes significantly reduced the Luc expression even compared to the RNP-treated group. This reduction in the Luc expression was further verified by immunofluorescence staining (Fig. S19), confirming decreased red Luc signal in the ternary complex-treated groups compared with non-treatment and RNP treatment groups. In addition, using the extracted and amplified target Luc DNA region, TIDE (Tracking of Indels by Decomposition) analysis was carried out to determine the indel frequency in the tumor samples (Fig. 6G). While RNP alone did not exhibit apparent indel frequency with only 1.5%, RNP/TA/P[Lys/Lys(FPBA)] showed 37.2% of the indel frequency (Fig. S20). RNP/TA/P[Glu(DET)/Glu(FPBA)] demonstrated the similar indel frequency with RNP/TA/P[Lys/Lys(FPBA)] (date not shown). Even there was no difference in the tumor volume (Fig. S21), RNP ternary complexes showed gene editing in the tumor sites. The Indel frequency was comparable in these ternary complexes, which is consistent with the luminescence measurement in Fig. 6D.

### Anti-tumor study

The therapeutic effect of RNP ternary complexes was examined using BALB/c nude mice bearing HeLa-Luc tumors (Figs 7A). We intravenously injected RNP samples (Cas9: 32.0 μg/mouse, sgRNA: 6.4 µg/mouse) containing polo like kinase 1 (PLK1) gene-targeted sgRNA 3 times, every other day (Table. S2). Naked RNP and RNP/TA complex did not show tumor growth inhibition effects due to the low accumulation and gene knockout efficacy in tumor sites. Similarly, RNP/TA/P[Lys/Lys(FPBA)] ternary complexes containing Luc gene-targeted sgRNA did not show tumor growth suppression (Fig. S21). Meanwhile, both RNP ternary complexes achieved the significant inhibition of the tumor growth as a result of the efficient *in vivo* tumor gene editing. Western blot and immunofluorescence staining further demonstrated the PLK1 knockout within the tumors (Fig. S22). According to the Western blot image, RNP/TA/P[Glu(DET)/Glu(FPBA)] therapy significantly reduced the PLK1 expression level, and RNP/TA/P[Lys/Lys(FPBA)] therapy also showed a tendency to decrease PLK1 expression compared to non-treated group and RNP treated group (Fig. S22A and B). Similarly, immunofluorescence staining observations revealed that both ternary complexes reduced the red PLK1 signal compared to PBS and RNP treated group (Fig. S22C), indicating *in vivo* PLK1 knockout function of RNP ternary complexes. TIDE analysis showed that ∼2% of Indels were introduced in the RNP ternary complexes (Fig. S23). In addition, treatment of the RNP ternary complexes exhibited similar weight changes and blood parameter values with treatment of D-PBS(-), indicating negligible adverse effects. (Figs 7B and S24, respectively).

**Figure. 7.**
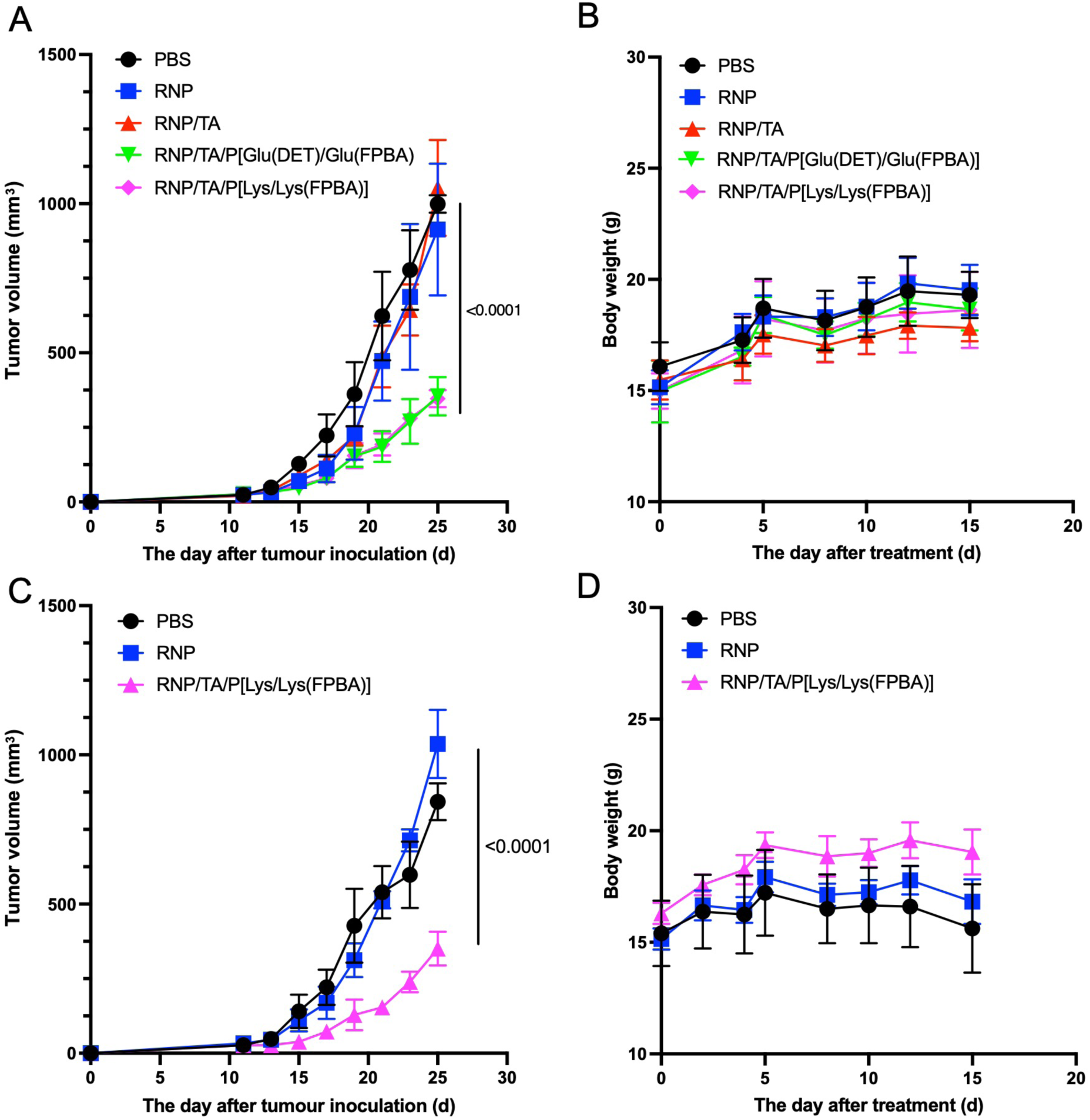
Anti-tumor study against HeLa-Luc and HCT116 subcutaneous tumor model mice. HeLa-Luc tumor cells were subcutaneously inoculated on day 0. PBS, RNP, RNP/TA, RNP/TA/P[Glu(DET)/Glu(FPBA)], RNP/TA/P[Lys/Lys(FPBA)] were administrated on day 11, day 13, and day 15. RNP was formed of Cas9 and sgRNA targeting PLK1. (A) The tumor growth curve and (B) body weight change of HeLa-Luc tumor model is shown. HCT116 tumor cells were subcutaneously inoculated on day 0, and PBS, RNP, RNP/TA/P[Lys/Lys(FPBA)] were administrated on day 11, day 13, and day 15. RNP was formed of Cas9 and sgRNA targeting KRAS^G13D^. (C) The tumor growth curve and (D) body weight change of HCT116 tumor model. The results are expressed as mean ± S.E.M. ANOVA with Tukey’s multiple comparison test was used for statistical analysis in both (A) and (C).

We then evaluated targeted gene editing efficacy using tumor cell line with a mutation in cancer related gene (Figs 7C and D). The mutation in Kirsten rat sarcoma virus (KRAS) promote malignancy of the cells, which is an undruggable target other than gene knockout technology. Especially, Human Colon Cancer (HCT) 116 cell line has a G13D mutation in KRAS gene ^32^. Using HCT 116 tumor bearing model mice, we also examined anti-tumor effect of RNP ternary complexes containing mutated KRAS-targeted sgRNA in the similar manner with the PLK1 study. The administration of RNP/TA/P[Lys/Lys(FPBA)] ternary complexes significantly impeded tumor growth compared to D-PBS(-) and RNP treatments, indicating the efficacy of this delivery system for precise gene editing in targeted cells, thus highlighting its therapeutic potential.

## 3. Discussion

The CRISPR/Cas system, derived from the bacterial immune system, has garnered significant attention as a transformative gene-editing tool. Gene-editing therapies hold tremendous promise for addressing various genetic disorders. To date, CRISPR/Cas-based therapies have been applied in clinical trials for congenital diseases, including Hemophilia and Mucopolysaccharidosis, typically *via* local injection. However, systemic application remains challenging due to limitations in site-specific targeting, insufficient stability in the bloodstream, and risks of off-target effects, thereby impeding clinical translation. While lipid-based nanoparticles have been primarily used for systemic genome editing strategies, their application has been limited mainly to the liver and spleen with a reticuloendothelial system^14,33,34^. There have been few reports of their use in other organs, such as tumors. In this study, we engineered a systemically applicable RNP delivery system utilizing PBA-conjugated polymers and tannic acid for tumor gene editing therapy. Most previous studies utilizing TA–PBA interactions in biomaterial-based drug delivery systems have primarily focused on the construction of pH-responsive hydrogels or applications involving local administration^35,36^. In contrast, the present study demonstrates the direct encapsulation of a large protein-nucleic acid complex within a well-defined supramolecular nanoscale assembly, enabling stimulus-responsive intracellular release and systemic *in vivo* tumor genome editing within a single polymer platform.

This ternary complex demonstrated exceptional tumor accumulation and effective gene-editing capabilities, showcasing its potential for cancer therapy. The RNP ternary complex is shielded by a PEG layer, which mitigates interactions with blood components and enhances circulatory stability. Its relatively large particle size compared to RNP, optimized for encapsulating a single RNP within the core, facilitates passive accumulation in tumor tissue via the enhanced permeability and retention (EPR) effect. In this biodistribution study, we observed that altering the side chains of the PBA-conjugated polymer influenced the biodistribution of the complex, suggesting the potential for organ-specific delivery. For instance, the P[Lys/Lys(FPBA)] polymer having primary amine side chain exhibited increased accumulation in the liver, whereas the P[Glu(DET)/Glu(FPBA)] polymer having ethylenediamine side chain predominantly localized in the kidney.

These findings highlight the importance of polymer structural modifications in modulating delivery profiles to minimize off-target effects and exhibit efficient gene editing in various tissues for treatment of genetic diseases. Our results demonstrated robust tumor-specific gene editing and significant tumor growth suppression without systemic toxicity. In the PLK1 and mutated KRAS editing study, the tumor growth was delayed even the knockout efficiency was a few percent. Still the kinetics of the editing efficiency in the tumor are under investigation, editing oncogene has a potential to treat the cancer. However, while tumor growth was arrested, complete remission was not observed in both anti-tumor study, indicating the inherent challenges of gene-editing-based cancer therapies. To further optimize efficacy, future efforts should focus on enhancing tumor accumulation and retention of the RNP complex. For example, modifying the PBA-conjugated polymer with tumor-targeting ligands, such as cyclic RGD peptides or glucose, could improve tumor specificity and accumulation^37^. Additionally, combining gene-editing therapies with other therapeutic modalities such as Olaparib, which is a type of PARP inhibitor against ovarian cancer, could yield synergistic effects^38^. In conclusion, our ternary complex system represents a promising strategy for tumor-specific gene delivery and editing, with significant potential to enhance current cancer therapies. By enabling the precise and effective delivery of active RNPs, this approach paves the way for a rational and integrative design of cancer treatment strategies.

## 4. Conclusion

In this study, we constructed systemic applicable RNP delivery systems by sequential self-assembly utilizing TA and PBA-conjugated polymers. This sequential self-assembled nanocarrier enhanced the tumor accumulation of RNP and demonstrated significant *in vivo* gene editing. In addition, RNP ternary complex armored with cancer gene targeting sgRNA introduced suppressed tumor growth in different tumor models. This RNP delivery system will offer as a systemic applicable cancer gene therapy.

## 5. Materials and Methods

Materials and methods to synthesize polymers were described in the Supplement information.

### Cells and animals

HeLa-Luc cell line was purchased from Caliper Life Science (Hopkinton, MA, USA). The cells were cultured in E-MEM medium containing 10% FBS and 1% penicillin-streptomycin in a humidified atmosphere containing 5% CO_2_ at 37 °C. HCT116 cell line was purchased from ATCC (Manassas, VA). The cells were cultured in McCoy’s 5a Medium with 10% Fetal bovine serum and 1% penicillin-streptomycin. Four-week-old female BALB/c nude mice were obtained from Japan SLC, Inc. (Hamamatsu, Japan). All the animal experiments were approved by the Animal Care and Use Committee of Tokyo Institute of Technology and performed in accordance with the Guidelines for the Care and Use of Laboratory Animals as stated by Tokyo Institute of Technology (2020-049).

### Preparation of Cas9 protein and sgRNA (RNP) complex

Cas9 protein (15 μg/μl) and sgRNA were mixed in the equal molar ratio in D-PBS(-) and incubated at room temperature for 10 min according to a manufacturer’s protocol of Synthego Co. Ltd. The sgRNA was used in a previous study for Luc gene editing^14^. The mixture solution was purified by ultrafiltration (MWCO: 10 kDa, Millipore, Burlington, MA, U.S.A.) against D-PBS(-) to remove the original solution containing glycerol. RNP complex comprising Cas9-GFP and sgRNA was prepared in the same manner. All RNP solution was diluted to specific concentration by D-PBS(-) or 10 mM HEPES for following experiments.

### Preparation of RNP/TA complexes

TA solution at various concentrations was prepared in D-PBS(-). The prepared TA solutions and RNP solutions (750 nM) were mixed at a volumetric ratio of 1:1 to prepare various molar ratios of TA to RNP ([TA]/[RNP] ratios). The prepared samples were incubated for 10 min at room temperature.

### Turbidity measurement of RNP/TA complex solution

Derived count rate of the RNP/TA complex solution (RNP: 750 nM) was measured as turbidity as previously reported using Zetasizer Nano ZS (Malvern Instruments, Worcestershire, UK) at a detection angle of 173°.

### Dynamic light scatter measurement

The hydrodynamic diameter of RNP samples (RNP: 750 nM) was measured in D-PBS(-) using a Zetasizer Ultra Red (Malvern Instruments) at a detection angle of 173° by a cumulant method. The measurement was performed at room temperature with a measurement time of 5 s and repeated 10 times.

### Preparation of RNP/TA/PBA-conjugated polymer ternary complexes

PBA-conjugated polymers (P[Glu(DET)/Glu(FPBA)] and P[Lys/Lys(FPBA)]) (1.0 mg/mL) aqueous solution was prepared in D-PBS(-). The RNP/TA complex ([TA]/[RNP]) = 30) was mixed with the PBA-conjugated polymer solution at the molar ratios of the polymer to RNP ([Polymer]/[RNP] ratios) of 30, followed by incubation for 10 min at room temperature. For *in vivo* experiment, the [Polymer]/[RNP] ratios of the ternary complex were set to 600.

### Size analysis using FCS

Hydrodynamic diameters of samples were calculated by fluorescence correlation spectroscopy (FCS) using an LSM710 confocal laser scanning microscope (CLSM, Carl Zeiss, Oberkochen, Germany) equipped with a ConfoCor module. An argon laser (488 nm) was used for excitation, and a 500−550 nm bandpass filter was used for detection. The prepared RNP samples incorporating Cas9-GFP in D-PBS(-) were put into an eight-well chamber with a concentration of 200 nM of Cas9-GFP. The measurement was performed at a room temperature with a sampling time of 10 sec, which was repeated 10 times. The hydrodynamic diameter was calculated from the obtained diffusion coefficients using the Stokes–Einstein equation with rhodamine 6G as a reference. Amplitude number of particles (AMPs) in the observation volume were also obtained in these measurements to calculate the number of encapsulated RNP molecules in complexes using the following equation:

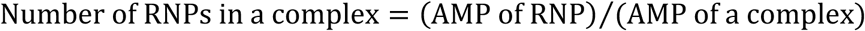

All measurements were conducted under identical optical alignment and fluorescence conditions to ensure consistent observation volume and detection efficiency. Note that, this calculation provides an average loading value, rather than indicating that every individual nanoparticle strictly contains one RNP molecule. To evaluate the stability of the complexes, D-PBS(-) containing FBS (10-50% FBS [v/v]) were utilized as the solvent. To evaluate pH-responsiveness of RNP ternary complexes, these hydrodynamic diameters were measured in D-PBS(-) (pH 7.4), 50 mM HEPES buffer (pH 6.8), and 50 mM 2-Morpholinoethanesulfonate (MES) buffer (pH 6.0 and 5.5).

### TEM observation

Three microliters of RNP samples (RNP: 1 mM) in 1 mM HEPES pH 7.4 were separately placed onto a copper grid coated with carbon film (mesh size 200 μm) and dried under vacuum. The grid was then set in a JEM-1400 TEM instrument (JEOL, Tokyo, Japan) for observation at an accelerating voltage of 100 kV.

### Agarose gel electrophoresis

RNP samples [1.1 μg of Cas9 for each sample in D-PBS(-)] were loaded in 1 % agarose gel and treated at 100 V for 20 min. The bands from RNP were visualized using CBB Stain One.

### FBS degradation stability

RNP samples (100 nM) were incubated with 0, 10, and 50 % FBS in PBS at 37 °C for 24 hours. The samples (4 μg of Cas9) were loaded in 1 % agarose gel and treated at 100 V for 20 min. The bands were visualized using SYBR® Green Ⅱ Nucleic Acid Gel Stain.

### RNase degradation stability

RNP samples (100 nM) were incubated with 1.0 mg/mL RNase at 37 °C for 2 hours. The samples (4 μg of Cas9) were loaded in 1 % agarose gel and treated at 100 V for 20 min. The bands were visualized using SYBR® Green Ⅱ Nucleic Acid Gel Stain.

### Apparent binding constants of PBA-conjugated polymers with GA

Apparent binding constants of PBA-conjugated polymers with GA were evaluated using ARS, following our previous study ^19,22^. The fluorescent intensity of the ARS/PBA-conjugated polymer complexes were measured with GA at various concentrations using a fluorophotometer (Spark, TECAN, Zürich, Switzerland). The apparent binding constants were calculated with the assumption that GA molecule should form a boronate ester with an FBPA molecule in polymers.

### Zeta-potential measurement of TA/PBA-conjugated polymer

Zeta-potential measurement of the TA/PBA-conjugated polymer complex (TA: 100 μM, PBA-conjugated polymer: 100 μM) in 10 mM HEPES was performed using a Zetasizer Nano ZS with a folded capillary zeta-potential cell (DTS1070, Malvern Instruments). The measurement was repeated 10 times.

### Subcellular distribution

HeLa-Luc cells were seeded on an eight-well chamber (1.0 × 10^4^ cells/dish) and incubated for 24 hours. After D-PBS(-) washing and medium replacement, the cells were incubated in the medium with RNP samples containing Cas9-GFP (RNP: 60 nM) for 24 hours. The cells were washed with D-PBS(-) and incubated for 30 min in D-PBS(-) containing 1 μM LysoTracker Red DND-99. Then, the cells were washed with D-PBS(-), and incubated for 3 min in D-PBS(-) containing 16.2 µM Hoechst 33342 (Life Technologies, Carlsbad, CA, USA) solution. The cells were washed with D-PBS(-), and observed in fresh D-PBS(-) using LSM710. Cas9-GFP, LysoTracker Red, and Hoechst 33342 were excited using laser light at 488 nm, 561 nm, and 405 nm, respectively. Subcellular distribution of RNP ternary complexes containing Cas9-GFP and Alexa647-labelled polymers were observed in the similar manner, and the Pearson correlation coefficient was calculated by the ZEN software (Carl Zeiss).

### Cellular uptake analysis

HeLa-Luc cells (1.0 × 10^4^ cells/well) were seeded into 96-well plates and incubated for 24 hours. The cells were washed with D-PBS(-) and incubated with RNP samples containing Cas9-GFP (RNP: 30 nM) in 100 µL of the cell culture medium for 24 hours. The cells were washed with D-PBS(-), treated with 50 μL of trypsin aqueous solution, and re-suspended with 150 μL of the cell culture medium. Cas9-GFP fluorescence intensities of the cells were measured using a flow cytometer (*Ex*/*Em* = 488/525 nm, Guava easy-Cyte 6-2 L, Merck Millipore, Billerica, MA, USA).

### *In vitro* Luc gene knockout study

HeLa-Luc cells (3.0 × 10^3^ cells/well) were seeded into 96-well plates and incubated for 24 hours. The cells were washed with D-PBS(-) and incubated with RNP samples (RNP: 30 nM) in 100 µL of the medium for 72 hours. After medium replacement, the cells were incubated for 1 hour in 110 μL of the cell culture medium with 10 % CCK-8 solutions. After the incubation, absorbance of the solution at 450 nm was measured using the fluorophotometer. The cells were washed with D-PBS(+) twice and incubated in 50 μL of cell culture lysis. After 15 min of incubation, 20 μL of the lysate was transferred to 96-well plate, and the luminescence intensity was measured using a Luciferase Assay System by GloMax 96 Microplate Luminometer (Promega Co. USA).

### Western blot to detect Luc expression *in vitro*

Hela-Luc cells treated with RNP samples (RNP: 30 nM) were washed with D-PBS(-) twice and incubated in 50 μL of cell culture lysis. 20 μg of protein was separated by sodium dodecyl sulfate-polyacrylamide gel electrophoresis (SDS-PAGE) using 4–15% Mini-PROTEAN® TGX™ Precast Protein Gels (Bio-Rad, Hercules, CA, USA) in Tris Glycine SDS (TGS) buffer. Then, the protein was transferred to a PVDF membrane. The membranes were blocked using 5 % BSA in TBS-T for 1 hour and washed with TBS-T. The membranes were incubated with anti-Firefly Luc antibody (1:1000, Abcam) or ß-actin antibody (1:6000, Abcam) dissolved in an immunoreaction enhancer solution (TOYOBO, Osaka, Japan) for overnight at 4 °C. After washing 3 times with TBS-T, horseradish peroxidase (HRP)-conjugated anti-rabbit antibodies (1:10000, Cell Signaling) were incubated as secondary antibodies. Membranes were visualized using SuperSignal™ West Dura Extended Duration Substrate (Thermo Fisher Scientific) and observed using an imaging system (Bio-Rad).

### *In vitro T7E1* study

HeLa-Luc cells (3.3 × 10^3^ cells/well) were seeded into 96-well plates and incubated for 24 hours. The cells were washed with D-PBS(-) and incubated with RNP samples (RNP: 30 nM) in 100 µL of the medium for 72 hours. After washing with D-PBS(-), 50 µL of trypsin-EDTA was added and incubated for 5 minutes to detach the cells. After washing the cells with D-PBS(-), the cells were resuspended with 50 µL of Cell Culture Lysis and incubated with 2 µL of Proteinase K following steps: 65°C for 15 min, 95°C for 10 min. The lysate was purified using QIAquick PCR Purification Kit (Qiagen) following the manufacture protocol. The targeted genomic loci were then amplified using the following PCR amplification program [94 °C for 30 s; (98 °C for 15 s; 66 °C for 25 s; 68 °C for 70 s) for 35 cycles; 68 °C for 30 sec and then keep at 4 °C]. The amplicons were then purified using QIAquick PCR purification kits and 400 ng of the purified DNA was added to 15 μL of annealing reaction containing 1 × NEBuffer 2. Then the PCR products were annealed in a thermocycler using the following conditions (95 °C for 5 min, then the mixture was cooled from 95 to 85 °C with Ramp Rate of -2 °C per second, following 85 to 25 °C with Ramp Rate of -0.1 °C per second, then keep at 4 °C) to form heteroduplex DNA. Afterwards, 1 μL of Resolvase (Takara Bio, Shiga, Japan) was added and incubated at 37 °C for 15 min. Next, the digested DNA was analysed using 1 % agarose gel electrophoresis and visualized using SYBR™ green I. All primers used for T7EI assay are listed in Table S3.

### 5.19. *In vitro* Luc DNA cleavage assay

The Luc target region DNA was extracted, following the protocol of *in vitro T7E1* study. The extracted Luc target region DNA with 300 ng was incubated with RNP samples (30 pmol) in D-PBS(-) pH 7.4-, or 50 mM MES buffer pH 5.5 for 60 min at [RNP]/[TA]/[PBA-conjugated Polymer] = 1/50/250. After the incubation, the reaction solution was mixed with 50mM Tris buffer (pH8.0) at 1/1(vol/vol) for neutralization. The mixture was isolated by agarose electrophoresis with DNA Ladder followed by SYBR safe staining to visualize DNA bands.

### Biodistribution of the RNP ternary complex

For evaluation of the biodistribution, fluorescently labelled sgRNA was prepared using Label IT® Cy5 Labeling kit (Mirus Bio LLC, Madison, WI, USA) according to the manufacturer’s protocol. The sgRNA and the labelling reagent were mixed at the molar ratio of 1.00 and 0.25, respectively. A subcutaneous HeLa-Luc tumor model was prepared by subcutaneously inoculating HeLa-Luc cells into the right back of BALB/c nude mouse (5.0 × 10^6^ cells/mouse). RNP samples containing Cy5 labelled sgRNA were intravenously injected to the mouse *via* teil vein at an RNP dose of 38.4 µg (32.0 µg of Cas9 and 6.4 µg of sgRNA)/mouse. Twenty μL of blood was collected from right teil vein at 1- and 10-hours post-injection. At 2-, 6-, and 24-hours post-injection, the mice were euthanized, and the organs were removed and washed using saline. The collected organs and blood were mixed with passive lysis buffer. After the homogenization and centrifugation at 10,000 g for 10 min at 4 °C, the supernatants were transferred to black 96 well plates, and the fluorescence of Cy5 was quantified using the fluorophotometer (*Ex*/*Em*: 650/667 nm, Spark).

### *In vivo* Luc gene knockout study

The subcutaneous HeLa-Luc tumor model was prepared by subcutaneously inoculating HeLa-Luc cells into the back of BALB/c nude mouse (4.0 × 10^6^ cells/mouse). When the tumor volume reached approximately 150 mm^3^, RNP samples were intravenously injected to the mouse *via* teil vein at an RNP dose of 38.4 µg/mouse. Luc activity on tumor was measured 13 minutes after intraperitoneal injection of luciferin using *in vivo* imaging system (IVIS, Perkin Elmer, Waltham, MA, USA) on day 0 before sample injections, and 3 days and 7 days after sample injections. At 7 days post-injection, the mice were sacrificed, and the tumors were collected and mixed with passive lysis buffer. After the homogenization, the luminescence intensities were measured using a Luciferase Assay System by Lumat LB9507 (Berthold Technologies, Bad Wildbad, Germany).

To detect the indels in the Luc gene, HeLa-Luc cells (4.5 × 10^6^ cells/mouse) were injected subcutaneously on the back flank of BALB/c nude mouse. When the tumor volume reached approximately 150 mm^3^, RNP samples were intravenously injected to the mouse *via* teil vein at an RNP dose of 38.4 µg/mouse on day 0, day 2 and day 4. The tumor tissue was harvested on day 16, and genomic DNA was extracted using DNeasy Blood & Tissue Kit following the manufacturer protocol. Then, the targeted gene sequence was amplified using the following PCR amplification program (98 °C for 2 min; (98 °C for 10 s; 60 °C for 15 s; 68 °C for 1 min) for 35 cycles; 68 °C for 30 sec and then keep at 4 °C). After purification, the amplicons were sent to the sanger sequencing (Reaction: Dye Terminator ver 3.1 (Applied Biosystems), Analysis: Genetic Analyzer 3730xl DNA Analyzer, (Thermo Fisher Scientific). The result was analysed using TIDE application.

### Western blot to detect Luc expression *in vivo*

RNP samples were intravenously injected to the HeLa-Luc tumor model mice *via* teil vein at an RNP dose of 38.4 µg/mouse on day 0, day 2 and day 4. The tumor tissue was harvested on day 16. The tumor was lysed using a lysis buffer containing protease/phosphatase inhibitors, and 20 μg of protein was separated by the SDS-PAGE. Subsequently, the protein was transferred onto PVDF membranes, and the membranes were treated in the same manner as for Western blot to observe Luc expression *in vitro*. The band intensity was calculated using the Image Lab Software, and the band intensity of Luc was normalized by that of ß-actin for densitometric quantification.

### Anti-tumor study

The subcutaneous HeLa-Luc tumor model or HCT116 tumor model were prepared by subcutaneously into the back of BALB/c nude mouse (4.5 × 10^6^ cells/mouse) on day 0. When the tumor volume reached approximately 30 mm^3^, RNP samples were intravenously injected to the mouse *via* teil vein at an RNP dose of 38.4 µg/mouse on day 11, -13 and -15. RNP was formed of Cas9 and sgRNA targeting PLK1/Luc for the treatment of HeLa-Luc tumor model mice, or KRAS^G13D^ for the treatment of HCT116 tumor model. The sgRNA designed to target KRAS^G13D^ incorporates the G13D mutation, specifically recognizing the mutated form of the KRAS gene (Table. S1). The tumor size was measured using a caliper, and the tumor volume (V) was calculated using the following equation: V = ab^2^/2, where a is the major axis, and b is a minor axis.

### Western blot to detect PLK1 expression *in vivo*

RNP samples were intravenously injected to the HeLa-Luc tumor model mice *via* teil vein at an RNP dose of 38.4 µg/mouse on day 0, day 2 and day 4. The tumor tissue was harvested on day 5. The tumor was treated in the same manner as the protocol for detecting Luc expression *in vivo* by Western blot. The extracted protein was separated by SDS-PAGE, followed by transferring onto a PVDF membrane. The membranes were blocked using 5 % BSA in TBS-T for 1 hour and washed with TBS-T. The membranes were incubated for overnight at 4 °C with anti-PLK1 antibody (1:2000, Abcam) or ß-actin antibody (1:6000, Abcam) dissolved in the immunoreaction enhancer solution. After washing 3 times with TBS-T, HRP-conjugated anti-rabbit antibodies were incubated as secondary antibodies. Membranes were visualized using SuperSignal™ West Dura Extended Duration Substrate and observed using the imaging system. The band intensity was calculated by the Image Lab Software, and the band intensity of PLK1 was normalized by that of ß-actin for densitometric quantification.

### Immunofluorescence staining

The collected tumor samples in the Western blotting study were fixed in p-formaldehyde overnight at 4 °C and then incubated in 15 and 30 wt% sucrose solution for each 2 h. The fixed tumor samples were sunk in OCT compound and sectioned at 5 μm. Cryosections were first incubated with 0.2 % Triton X-100 for 15 minutes, then blocked with 2% BSA in PBS at room temperature for 1 hour. Tissue samples were stained with rabbit anti-Luc antibodies (1:100, Abcam) or anti-PLK1 antibodies (1:100, Abcam) for 2 hours at room temperature. After washing the slides with PBS-T 3 times, tissue samples were then stained with the Alexa-647-labelled anti-rabbit IgG (1:500, Cell Signaling) for 1 hour at room temperature. After washing the slides with PBS-T 3 times, sections were then covered with Vector TrueVIEW® Autofluorescence Quenching Kit with DAPI (Vector Laboratories, Inc., SP-8500). Sections were observed under a BZ-X800 microscope (KEYENCE, Tokyo, Japan).

### Toxicity study

RNP samples containing PLK1 targeting sgRNA were intravenously injected to female 7-week-old BALB/c nude mice *via* teil vein at an RNP dose of 38.4 µg/mouse on day 0, - 3, and -6. Seven days post treatment, the blood was collected, and plasma was collected by centrifuging the blood (5000 rpm, 10 min). The WBC (white blood cells), RBC (red blood cells), PLT (platelet) in blood was analysed an Automatic Multiple Blood Cell Counter (POCH-100IVDIFF, Sysmex, Lincoln-shire, IL, U.K.). BUN (blood urea nitrogen), CRE (creatinine), GPT-P (glutamic pyruvic transaminase), GOT/AST (Glutamic Oxaloacetic Transaminase/Aspartate transaminase), and LDH (lactic acid dehydrogenase) levels were analysed in the plasma using FujiDri-Chem (Fujifilm, Tokyo, Japan).

### Statistical analysis

Statistical significances (*P < 0.05, **P < 0.01, ***P < 0.001) were determined using Student’s t-test for analyzing two groups. For multiple comparisons, one-way ANOVAs with Turkey’s test was used.

## Acknowledgments

We thank the Division of Materials Analysis Suzukake-dai, Technical Department, Institute of Science Tokyo, for ^1^H NMR analysis. We would like to thank Editage (www.editage.com) for English language editing.

## Author Contributions

T.M. and Y.H. designed and performed all experiments. T.M. and Y.H. wrote the manuscript. T.C. assisted *in vivo* experiments. T.N. and Y.O. advised *in vitro* and *in vivo* experiments and proofread the manuscript. Y.M. advised polymerization. N.N. and Y.H. supervised the whole project and conceived the concept of this study. This manuscript was written through contributions of all authors. All authors have given approval to the final version of the manuscript.

## Data Availability Statement

The data that support the findings of this study are available from the corresponding author upon reasonable request.

## Supporting Information

Materials; Synthesis of PEG-Plys.; Synthesis of PEG-P[Glu(DET)].; Synthesis of PBA-conjugated polymers.; Fluorescence labelling; Turbidity measurement by light scattering analysis. RNP concentration: (750 nM); Synthetic scheme of PEG-P[Lys/Lys(FPBA)]; Synthetic scheme of PEG-P[Glu(DET)/Glu(FPBA)]; GPC chart of PEG-PBLG and PEG-P[Glu(DET)]; ^1^H-NMR spectra for PEG-PBLG; ^1^H-NMR spectra of PEG-PEG-P[Glu(DET)] and PEG-P[Glu(DET)/Glu(FPBA)]; ^1^H-NMR spectra of PEG-P[Lys/Lys(FPBA)]; Size and PDI measurement using DLS; TEM images of RNP samples; RNase degradation stability; pH-dependent hydrodynamic diameters of RNP/ ternary complexes; Large scale subcellular distribution observed by CLSM; Intracellular dissociation of the RNP ternary complex by CLSM; Representative western blot images *in vitro*; *In vitro* DNA cleavage assay.; Zeta potential of TA samples; Tumor size of subcutaneous Hela-Luc after RNP sample containing Luc-target sgRNA-treatment; Immunofluorescence staining images of Hela-Luc tumors treated with RNP samples targeting Luc; TIDE analysis of sgLuc target loci; The tumor growth curve of Hela-Luc tumor model; TIDE analysis of sgPLK1 target loci; *In vivo* PLK1 expression level in Hela-Luc tumors-treated with RNP samples; Biochemical parameters of blood; Sequences of sgRNA used in this study; *Mw/Mn* of PEG-P[Lys(TFA)] and PEG-PBLG; Sequences of primers used in this study.

## ABBREVIATIONS

CRISPR: Clustered regularly interspaced short palindromic repeats;
Cas9: CRISPR-associated protein 9;
sgRNA: single guide RNA,
RNP: ribonucleoprotein.
TA: tannic acid;
PBA: phenylboronic acid;
PEG: poly(ethylene glycol);
PEG-PBLG: PEG–*b*-poly(γ-benzyl-L-glutamate);
PEG-PLys: PEG-*b*-poly(L-lysine);
DET: diethylenetriamine;
FPBA: 4-carboxy-3-fluorophenylboronic acid;
PBA: Phenylboronic acid;
KRAS: Kirsten rat sarcoma virus;
PLK1: polo-like kinase 1;
DLS: dynamic light scattering;
FCS: fluorescence correlation spectroscopy;
ARS: Alizarin red S;
MES: 2-Morpholinoethanesulfonate;
GFP: green fluorescent protein;
CLSM: Confocal Laser Scanning Microscopy;
T7EI: T7 Endonuclease I;
Luc: luciferase;
IVIS: *in vivo* imaging system;
TIDE: Tracking of Indels by Decomposition;
TBS-T: Tris-Buffered saline with Tween 20;
HRP: Horseradish peroxidase;
PVDF: Polyvinylidene difluoride;
SDS-PAGE: Sodium dodecyl sulfate–polyacrylamide gel electrophoresis;
BSA: bovine serum albumin;
PFA: Paraformaldehyde;
PDI: Protein disulfide isomerase;
TEM: Transmission electron microscopy.

Title: CRISPR-Cas9 RNP-loaded ternary complexes composed of tannic acid and phenylboronic acid-conjugated polymers significantly enhance systemic stability, enabling prolonged blood circulation and efficient tumor accumulation compared to RNP. Following endocytosis and endosomal escape, localized RNPs perform gene editing and selectively eliminate cancer cells, resulting in potent therapeutic efficacy across multiple tumor models.

## ToC figure

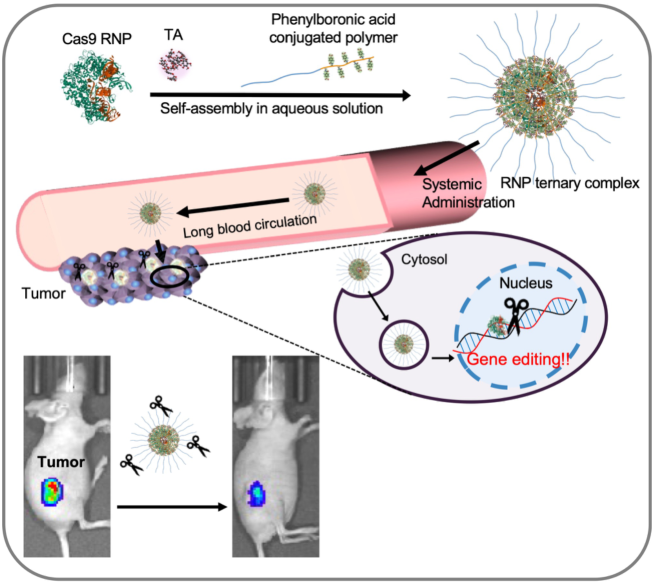

