## Supplementary Figures for "Sequentially Self-Assembled Supramolecular Nanocomplexes Enable Systemic Cas9 RNP Delivery and *In Vivo* Tumor Genome Editing"

This PDF file includes:
Supplementary Materials.
Supplementary Method. Synthesis of PEG-Plys.
Supplementary Method. Synthesis of PEG-P[Glu(DET)].
Supplementary Method. Synthesis of PBA-conjugated polymers.
Supplementary Method. Fluorescence labelling.
Supplementary Figure. S1. Turbidity measurement by light scattering analysis.
Supplementary Figure. S2. Synthetic scheme of PEG-P[Lys/Lys(FPBA)].
Supplementary Figure. S3. Synthetic scheme of PEG-P[Glu(DET)/Glu(FPBA)].
Supplementary Figure. S4. GPC chart of PEG-PBLG and PEG-P[Glu(DET)].
Supplementary Figure. S5. $^1$H-NMR spectrum of PEG-PBLG.
Supplementary Figure. S6. $^1$H-NMR spectrum of PEG-P[Glu(DET)] and PEG-P[Glu(DET)/Glu(FPBA)].
Supplementary Figure. S7. $^1$H-NMR spectrum of PEG-P[Lys/Lys(FPBA)].
Supplementary Figure. S8. Size and PDI measurement using DLS.
Supplementary Figure. S9. TEM images of RNP samples.
Supplementary Figure. S10. Serum degradation stability
Supplementary Figure. S11. RNase degradation stability.
Supplementary Figure. S12. pH-dependent hydrodynamic diameters of RNP/ ternary complexes.
Supplementary Figure. S13. Large scale subcellular distribution observed by CLSM
Supplementary Figure. S14. Intracellular dissociation of the RNP ternary complex by CLSM.
Supplementary Figure. S15. Representative Western blot images *in vitro*.
Supplementary Figure. S16. *In vitro* DNA cleavage assay.
Supplementary Figure. S17. Zeta potential of TA samples.
Supplementary Figure. S18. Tumor size of subcutaneous Hela-Luc after RNP sample containing Luc-target sgRNA-treatment.
Supplementary Figure. S19. Immunofluorescence staining images of Hela-Luc tumors treated with RNP samples targeting Luc.
Supplementary Figure. S20. TIDE analysis of sgLuc target loci.
Supplementary Figure. S21. The tumor growth curve of Hela-Luc tumor model using control sgRNA.

Supplementary Figure. S22. *In vivo* PLK1 expression level in Hela-Luc tumors-treated with RNP samples.

Supplementary Figure. S23. TIDE analysis of sgPLK1 target loci.

Supplementary Figure. S24. Biochemical parameters of blood.

Supplementary Table. S1. Polymer information.

Supplementary Table. S2. sgRNA sequence.

Supplementary Table. S3. Primer sequence.

### Supplemental methods

#### Materials

α-Methoxy-ω-propylamine-poly(ethylene glycol) (MeO-PEG-$NH_2$) ($M_n$:10,000) was purchased from NOF corporation (Tokyo, Japan). ε-Trifluoro acetyl-L-lysine *N*-carboxy anhydride (Lys(TFA)-NCA), γ-Benzyl-L-glutamic Acid *N*-carboxy anhydride (BLG-NCA) was purchased from Chuo Kaseihin Co., Inc. (Tokyo, Japan). Alizarin red S (ARS), gallic acid (GA), TA, benzene, hydrochloric acid (HCl), sodium hydroxide solution (NaOHaq), Sodium Acetate, Eagle's Minimum Essential Medium with L-Glutamine and Phenol Red (E-MEM), boric acid, and 4-(4,6-dimethoxy-1,3,5-triazin-2-yl)-4-methylmorpholinium chloride (DMT-MM) were purchased from FUJIFILM Wako Pure Chemical Industries, Ltd. (Osaka, Japan). Sodium hydrogen carbonate ($NaHCO_3$), D-sorbitol, Diethylenetriamine (DET), 2-hydroxypyridine, and Tween 20 were purchased from Tokyo Chemical Industry Co., Ltd. (Tokyo, Japan). Diethyl ether, dimethyl sulfoxide-$d_6$ + 0.05% V/V TMS (D, 99.9%) (d-DMSO) were purchased from Kanto Chemical Co., Inc. (Tokyo, Japan). Methanol (MeOH), Dulbecco's PBS(-) [D-PBS(-)], dimethyl sulfoxide (DMSO), CBB Stain One, Tris(hydroxymethyl) aminomethane (Tris), and sodium chloride (NaCl) were obtained from Nacalai Tesque, Inc. (Kyoto, Japan). 1-Methyl-2-pyrrolidinone (NMP), lithium bromide (LiBr), Deuterium oxide, sodium chloride, trypsin/EDTA, penicillin-streptomycin solution, Triton X-100, Agarose S and Cas9-GFP Protein were purchased from Sigma-Aldrich Co. (St. Louis, MO, U.S.A.). Modified sgRNAs (Table S1) were purchased from Synthego (Menlo Park, CA, USA). LysoTracker Red DND-99, Hoechst 33342, Alexa647-NHS, SuperSignal™ West Dura Extended Duration Substrate, and DNase/RNase-Free Distilled Water were purchased from Thermo Fisher Scientific (Waltham, MA, USA). Passive lysis buffer, Cell culture Lysis, Luciferin (In Vivo Grade), and Luciferase Assay System were purchased from Promega (Madison, WI, USA). 4-(2-Hydroxyethyl)-1-piperazineethanesulfonic acid (HEPES), 2-Morpholinoethanesulfonate (MES), 2NA (EDTA•2Na), and Cell Counting Kit-8 were purchased from Dojindo (Kumamoto, Japan). Cas9 NLS Nuclease (15 μg/μl) and 6 x Loading Buffer Triple Dye were purchased from NIPPON GENE CO., LTD. (Tokyo, Japan). Guide-it™ Mutation Detection Kit, SYBR® Green I Nucleic Acid Gel Stain and SYBR® Green II Nucleic Acid Gel Stain were purchased from Takara Bio Inc. (Shiga, Japan). Cell culture dishes and glass base dishes were obtained from AGC Techno Glass Co., Ltd. (Shizuoka, Japan). Eight-well chamber was purchased from Nalge Nunc International (NY, USA). Syringe filters were obtained from ADVANTEC TOYO KAISHA, LTD. (Tokyo, Japan). Fetal Bovine Serum (FBS) was purchased from Biosera Co., Ltd. (Caille, France). Proteinase K, DNeasy Blood & Tissue Kit, and QIAquick PCR Purification Kit were purchased from Qiagen (Hilden, Germany). Albumin, Bovine Serum, General Grade, pH7.0 (BSA) was purchased from Nacalai Tesque. Inc (Tokyo, Japan). 4–15% Mini-PROTEAN® TGX™ Precast Protein Gels, 10x Tris/Glycine/SDS, and Trans-Blot Turbo Mini 0.2 μm PVDF Transfer Packs were purchased from Bio-Rad (Hercules, CA, USA). Rabbit anti-Firefly Luciferase antibody, Rabbit anti-PLK1 antibody, and Rabbit anti-β-actin antibody were purchased from Abcam. Inc (MA, USA). Anti-rabbit IgG (H+L), F(ab')2 Fragment (Alexa Fluor® 647 Conjugate) and Anti-rabbit IgG, HRP-linked Antibody were purchased from Cell Signaling (Leiden, Netherland). Vector TrueVIEW® Autofluorescence Quenching Kit with DAPI was purchased from Vector Laboratories, Inc. (CA, USA).

**Synthesis of PEG-PLys**

PEG-PLys was synthesized as previously reported (Fig. S2)[1]. Briefly, MeO-PEG-$NH_2$ was lyophilized with benzene and dissolved in distilled DMSO. The solution was mixed with distilled DMSO containing Lys(TFA)-NCA and stirred at room temperature under argon atmosphere. After 3 days reaction, the mixture was poured into an excess amount of diethyl ether to obtain white precipitates of PEG-poly(ε-trifluoro acetyl-L-lysine) (PEG-P[Lys(TFA)]). The obtained polymer was characterized by gel permeation chromatography (GPC) using an HPLC system (JASCO, Tokyo, Japan) equipped with two TSKgel superAW3000 and superAW4000 (Tosoh Corporation, Tokyo, Japan) in NMP containing LiBr (50 mM) at a flow rate of 0.3 mL/min at 40 °C. The $M_w/M_n$ of the polymer was calculated to be 1.16 based on PEG calibration (Table S2). To deprotect the trifluoro acetyl group, PEG-PLys(TFA) was dissolved in a mixture of methanol/5 M NaOH [4:1 (v/v)], and stirred at room temperature for 24 hours. The mixture was dialyzed against 5mM HCl aqueous solutions and subsequently against deionized water. Then, the dialyzed solution was lyophilized to obtain PEG-P[Lys] as a white powder. PEG-P[Lys] was characterized in $D_2O$ using $^1$H NMR (AVANCE III 400, Bruker, Billerica, MA, USA), and the DP of Lys unit was estimated to be 20 by comparing the ratio of peak area in butylene protons in P[Lys] side chain and oxyethylene protons in PEG.

**Synthesis of PEG-P[Glu(DET)]**

PEG-P[Glu(DET)] was synthesized as previously reported (Fig. S2).[2] For synthesis of poly(γ-benzyl-L-glutamate) (PEG-PBLG), MeO-PEG-$NH_2$ solution in DMSO was mixed with DMSO containing BLG-NCA and stirred at room temperature under argon atmosphere for 3 days. The mixture was poured into diethyl ether to obtain a white precipitate of PEG-PBLG. The obtained polymer was characterized by GPC in the same manner as the characterization of PEG-PLys(TFA) (Fig. S4A). The $M_w/M_n$ was calculated to be 1.08 based on PEG calibration (Table S2). The DP of Glu was determined to be 24 from the peak intensity ratio of the ethylene protons in PGlu side chain to the protons derived from oxyethylene in PEG by $^1$H NMR spectroscopy (Fig. S5). To modify the side chain of the PBLG segment with diethylenetriamine (DET), aminolysis was performed. Briefly, lyophilized PEG-PBLG was dissolved in NMP. In addition, 2-hydroxypyridine (25 eq. to Glu unit) and DET (250 eq. to Glu unit) were dissolved in NMP, separately. Then, these solutions were mixed under argon atmosphere and stirred for 3 days at room temperature. The mixture solution was dialyzed against 0.01 M HCl aqueous solution and subsequently against deionized water. The dialyzed solution was then lyophilized to obtain PEG-P[Glu(DET)] as a white powder. The obtained PEG-P[Glu(DET)] was characterized by $^1$H-NMR spectroscopy (Fig. S6A) and GPC [column: Superdex 200 Increase 10/300 GL, eluent: 10 mM HEPES with 500 mM NaCl, flow rate: 0.75 mL/min, detection: absorbance at 220 nm] (Fig. S4B).

**Synthesis of PBA-conjugated polymers**

The synthesis was performed according to the previous reported study (Figs. S1 and S2)[1]. FPBA (50 % eq. of Lys unit) was introduced into the side chains of PEG-P[Lys] (1eq.) by a condensation reaction using DMT-MM in methanol containing 50 mM $NaHCO_3$ solution (pH 8.5) for 24 hours at room temperature. After the reaction, the mixture was dialyzed against 5 mM HCl aqueous solutions and subsequently against deionized water. Then, the dialyzed solution was lyophilized to obtain PEG-P[Lys/Lys(FPBA)], termed P[Lys/Lys(FPBA)], as a white powder. The rate of FPBA modification was determined to be 50% from the peak intensity ratio of the propylene protons in PLys side chain to the protons derived from the FPBA group by $^1$H-NMR spectroscopy using $D_2O$ containing 50 mg/mL of D-sorbitol (Fig. S7). FPBA-modified PEG-P[Glu(DET)], which were termed P[Glu(DET)/Glu(FPBA)], were synthesized and characterized in the similar manner as the preparation of P[Lys/Lys(FPBA)]. The number of conjugated FPBAs were calculated to be 10 and 14 on P[Lys/Lys (FPBA)] and P[Glu(DET)/Glu(FPBA)], respectively, by $^1$H-NMR spectroscopy (Fig. S6B).

**Fluorescence labelling**

P[Lys/Lys (FPBA)] and P[Glu(DET)/Glu(FPBA)] were mixed with Alexa647-NHS (equivalent moles of the polymer) overnight in 10 mM $NaHCO_3$ and 140 mM NaCl buffer (pH 7.4). The mixture was purified by dialysis with DI water twice (MWCO: 3 kDa) and PD-10 desalting columns packed with Sephadex™ G-25 resin (Cytiva, Marlborough, USA) following the manufacture's protocol. The labeled polymers were obtained by freeze-drying as powders.

Supplemental figures

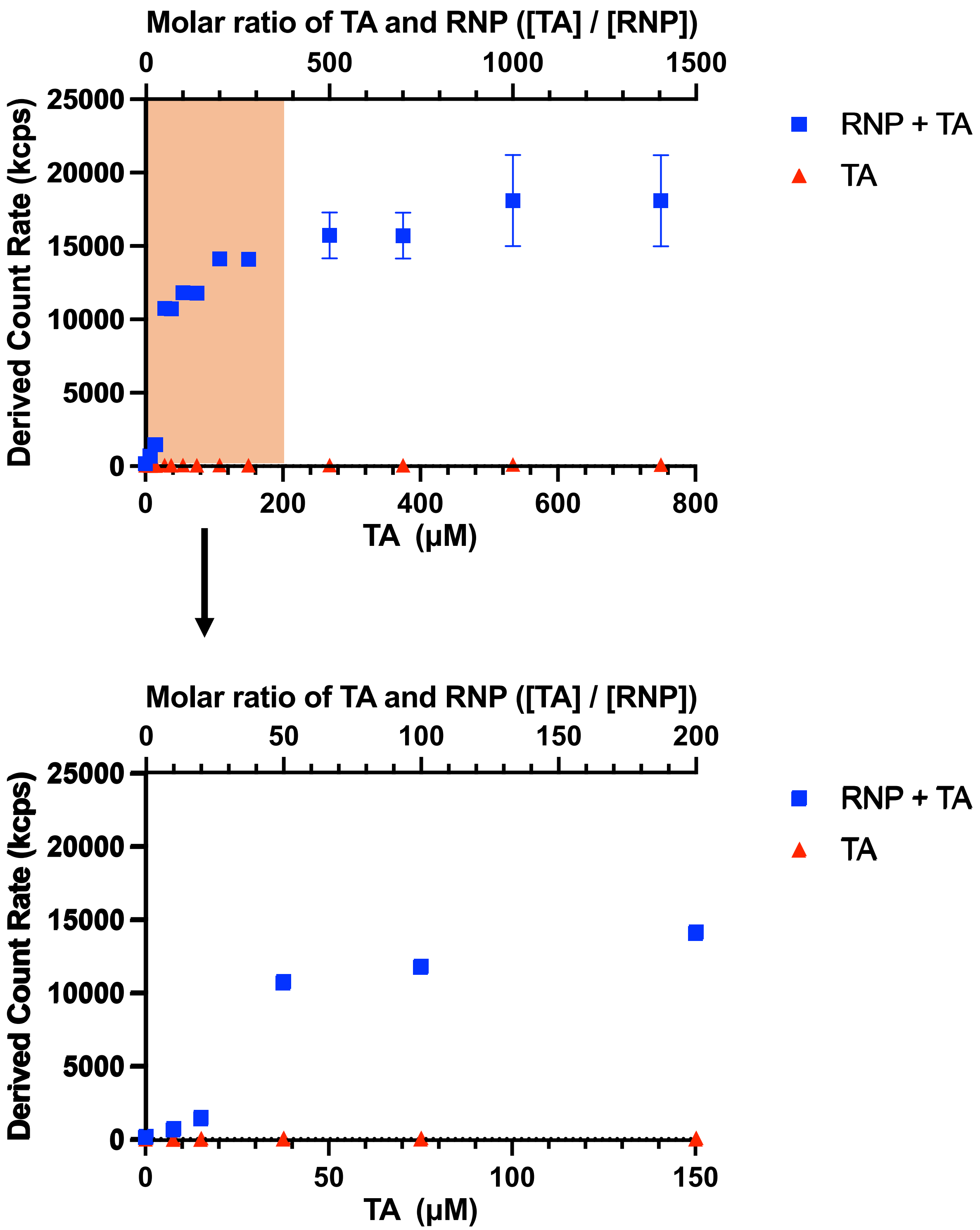


**Fig. S1.** Turbidity measurement by light scattering analysis. RNP concentration: (750 nM).

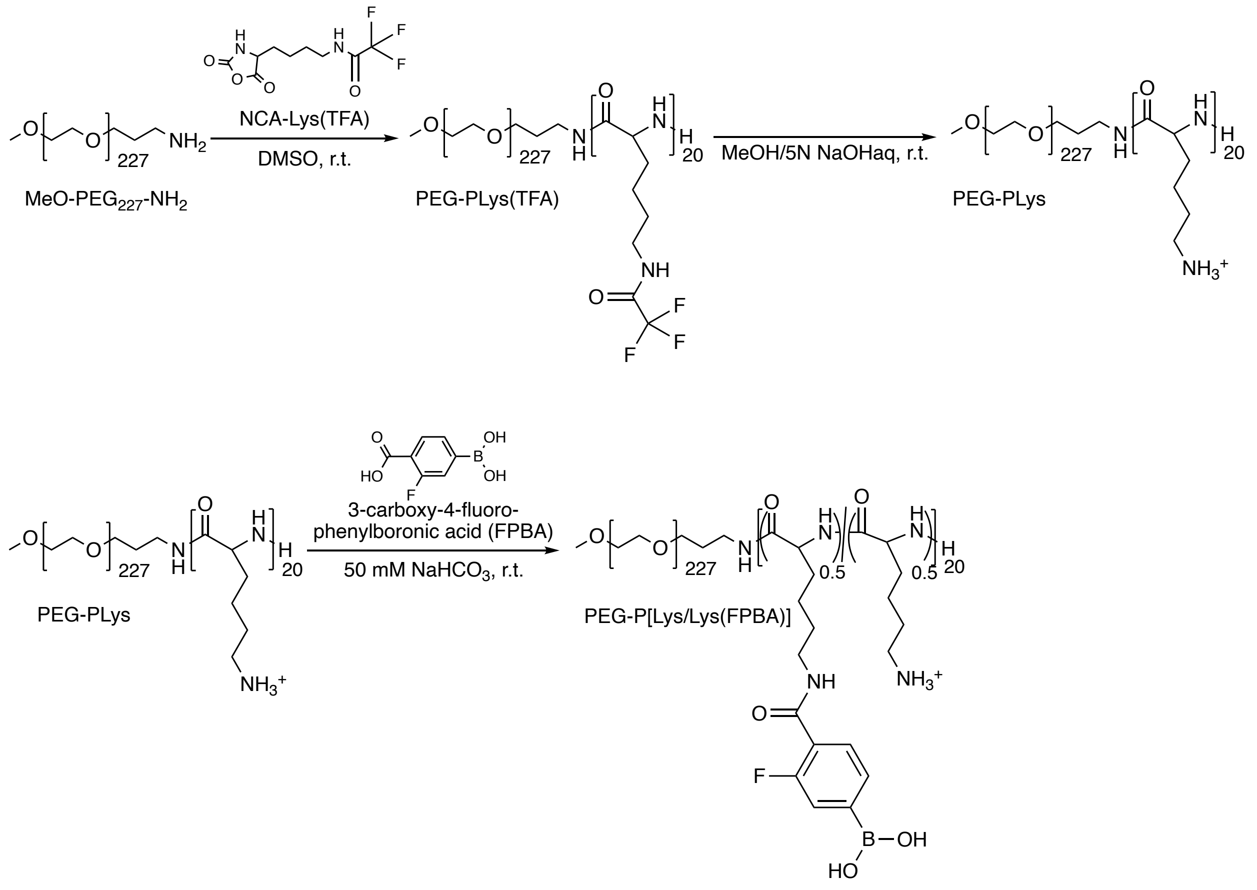


**Fig. S2.** Synthetic scheme of PEG-P[Lys/Lys(FPBA)].

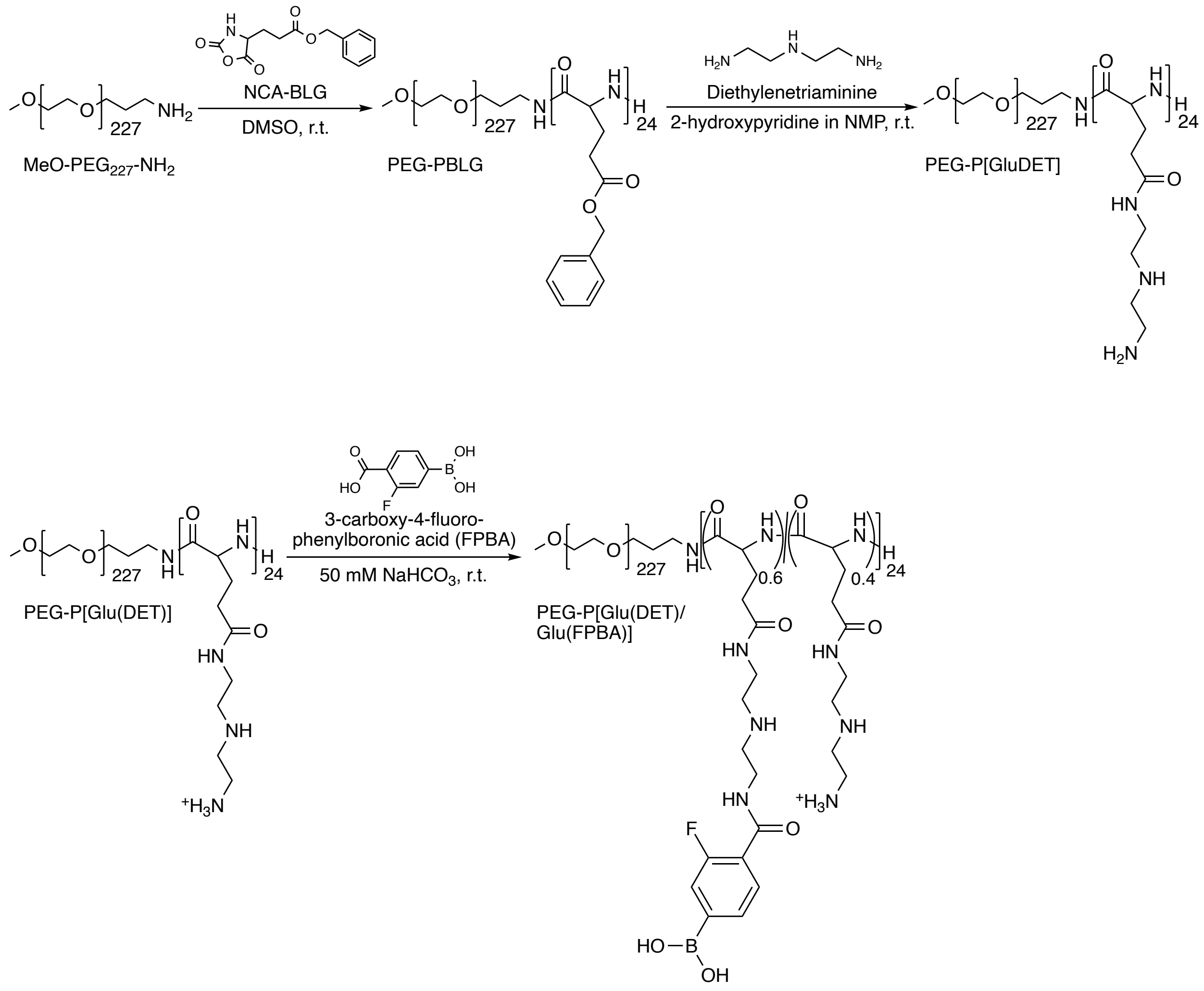


**Fig. S3.** Synthetic scheme of PEG-P[Glu(DET)/Glu(FPBA)].

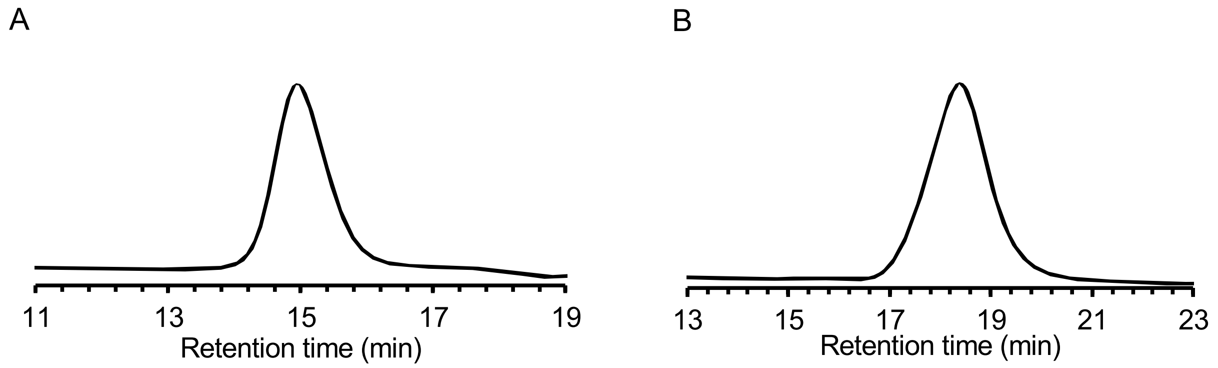


**Fig. S4.** (A) GPC chart of PEG-PBLG. Column: two TSKgel superAW3000 and superAW4000 (Tosoh Corporation, Tokyo, Japan), detector: a refractive index detector, eluent: NMP containing LiBr (50 mM), flow rate: 0.2 mL/min, temperature: 40 °C. (B) GPC chart of PEG-P[Glu(DET)]. Column: Superdex 200 Increase 10/300 GL, eluent: 10 mM HEPES buffer (pH 7.4) containing 500 mM NaCl, flow rate: 0.75 mL/min, temperature: room temperature, detection: absorbance at 220 nm.

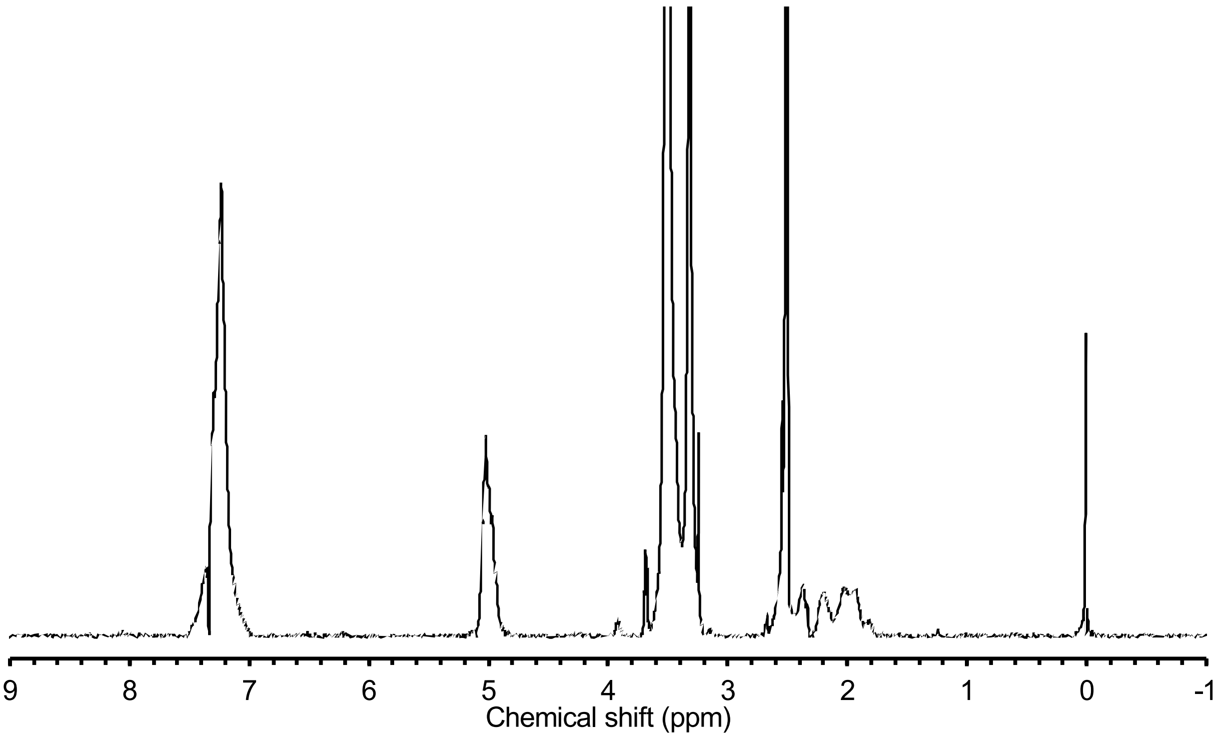


**Fig. S5.** $^{1}$H-NMR spectrum of PEG-PBLG. (solvent: DMSO-$d_6$).

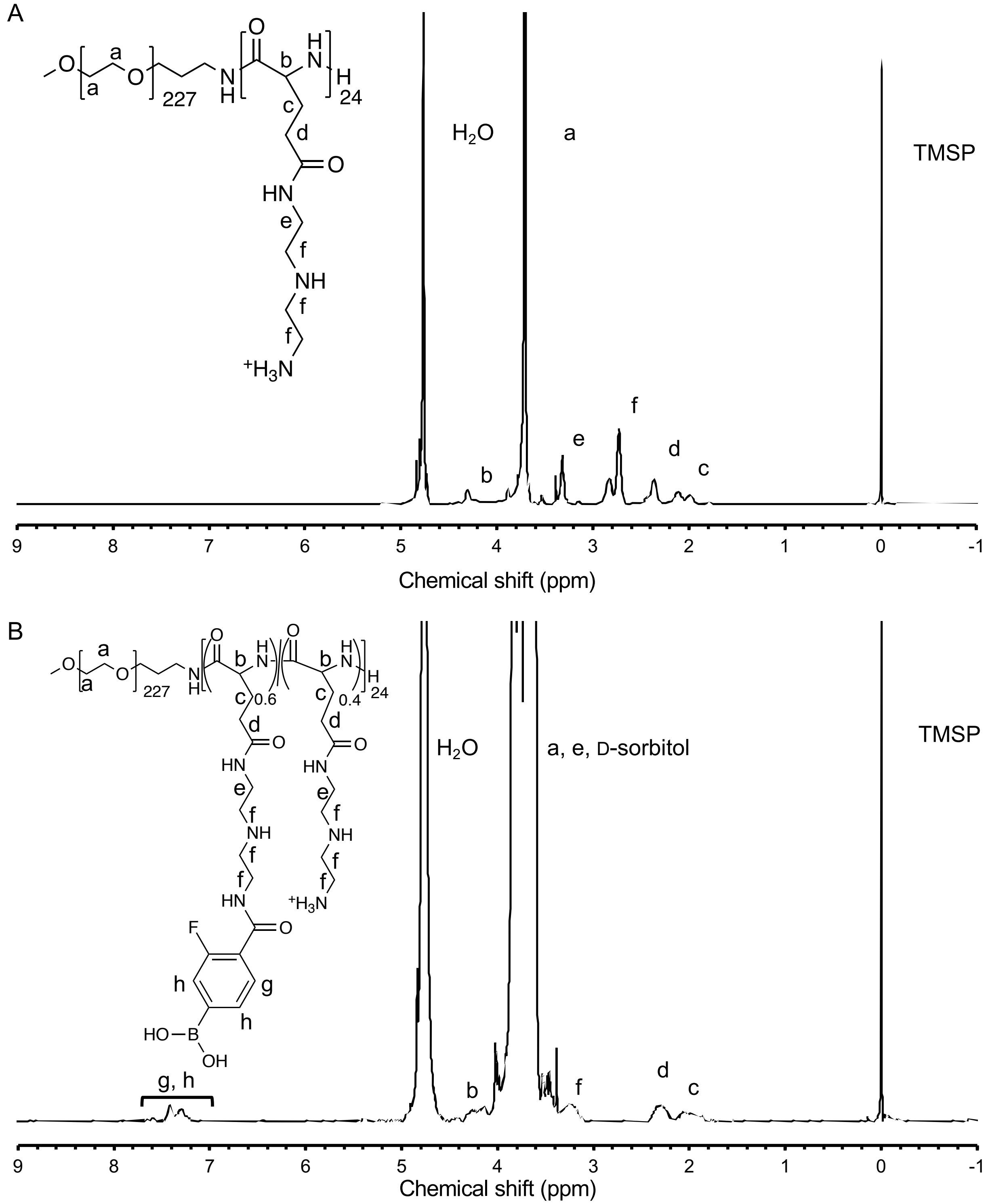


**Fig. S6.** ¹H-NMR spectrum of (A) PEG-P[Glu(DET)] (solvent; $D_2O$) and (B) PEG-P[Glu(DET)/Glu(FPBA)] (solvent: $D_2O$ with 50 mg/mL D-sorbitol).

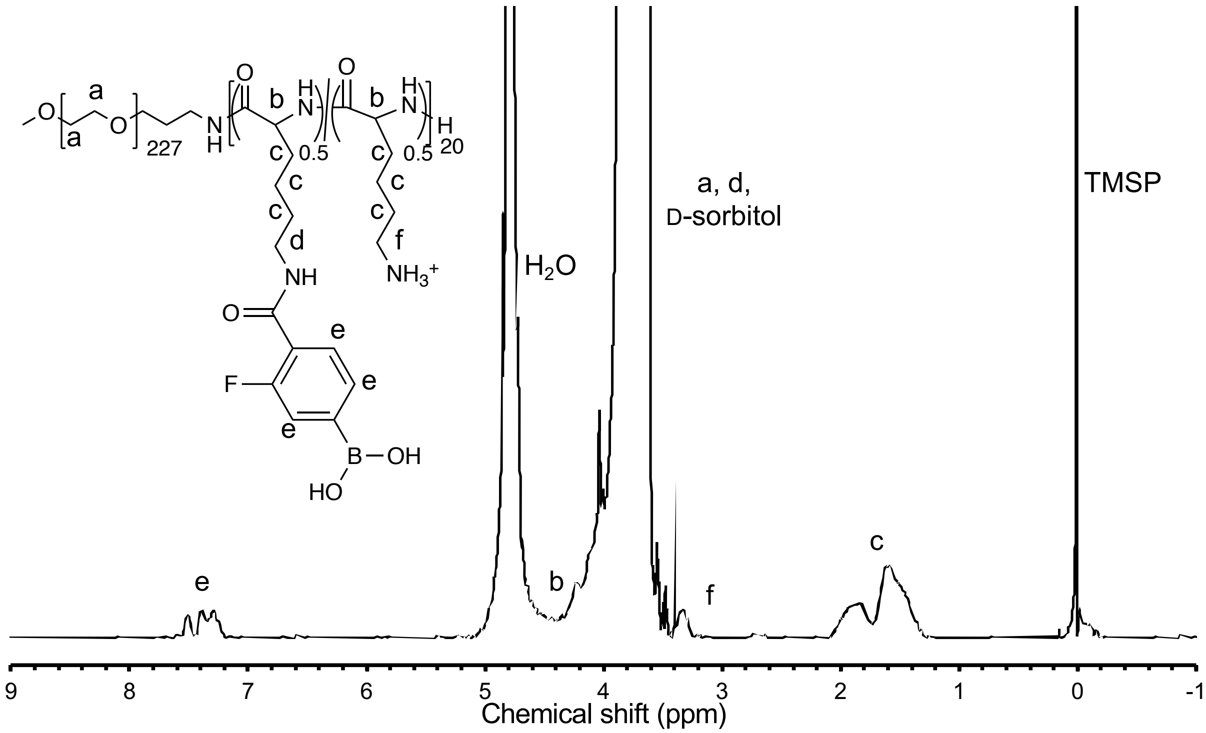


**Fig. S7.** $^1$H-NMR spectrum of PEG-P[Lys/Lys(FPBA)] (solvent: $D_2O$ with 50 mg/mL D-sorbitol).

A

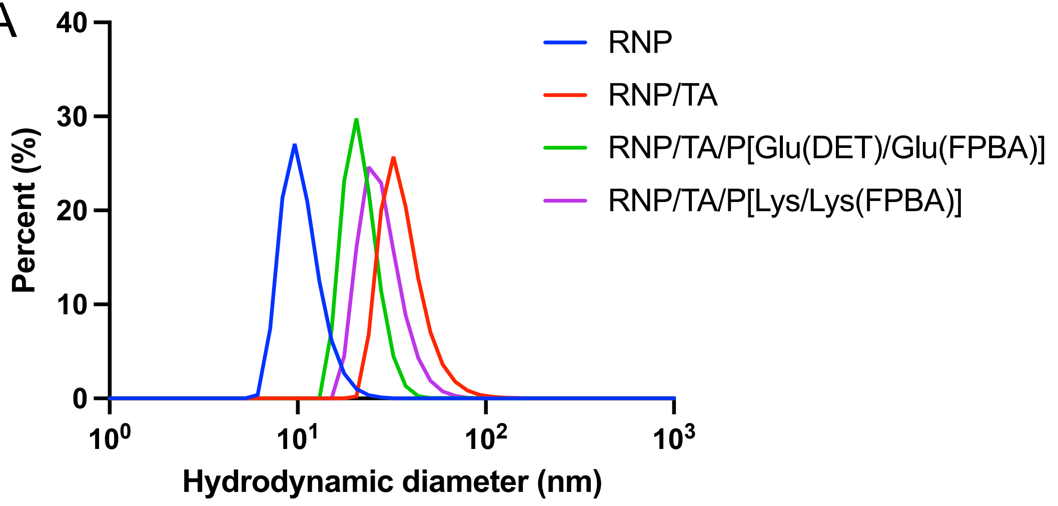


B

| Sample | RNP | RNP/TA | RNP/TA/ P[Glu(DET) /Glu(FPBA)] | RNP/TA/ P[Lys /Lys(FPBA) |
|---|---|---|---|---|
| PDI ± S.D. | 0.35 ± 0.02 | 0.31 ± 0.03 | 0.13 ± 0.02 | 0.29 ± 0.01 |

**Fig. S8.** Size and PDI measurement using DLS. (A) Size measurement and (B)PDI of RNP samples. The results are expressed as mean ± S.D. (n = 10).

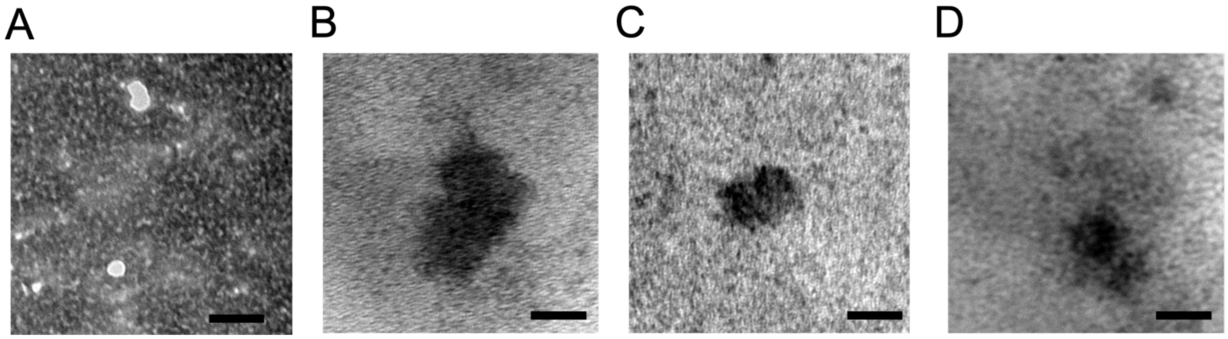


**Fig. S9**. TEM images of RNP samples. (A) RNP, (B) RNP/TA, (C) RNP/TA/P[Glu(DET)/Glu(FPBA)], and (d) RNP/TA/P[Lys/Lys(FPBA)] ternary complex in air condition. Scale bar: 20 nm.

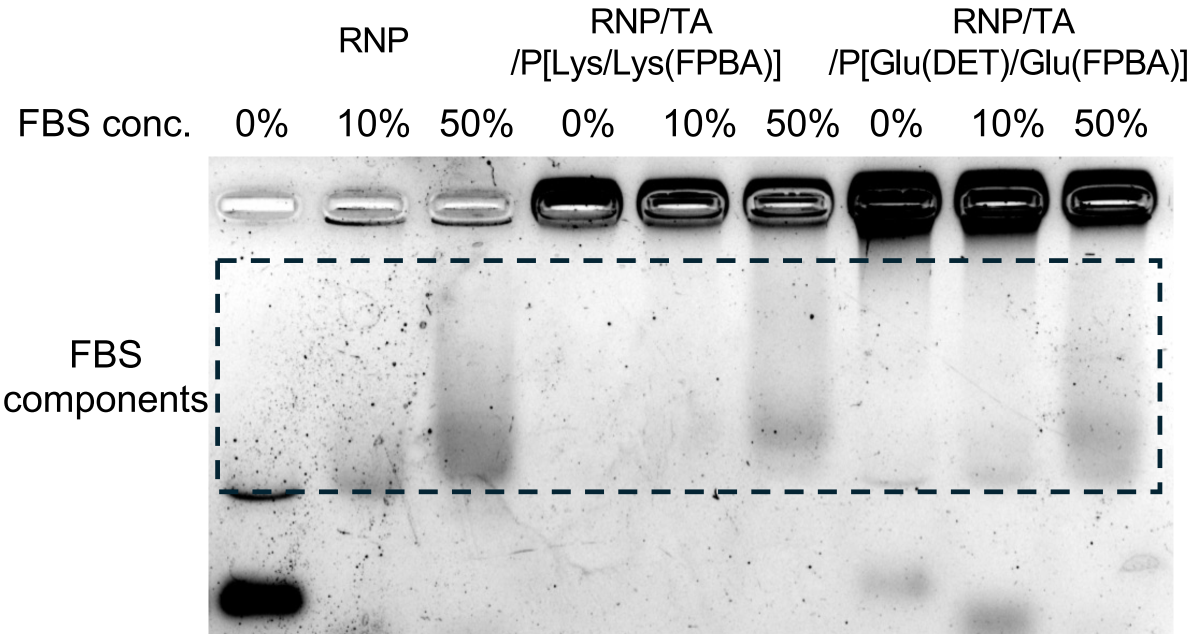


**sFig. S10**. Serum degradation stability detected by 1 % agarose gel electrophoresis. RNP concentration: 1 μM, [TA]/[RNP] = 30, [PBA-conjugated polymer]/[RNP] = 60. RNP samples were incubated with 10 or 50 % of FBS in aqueous solution.

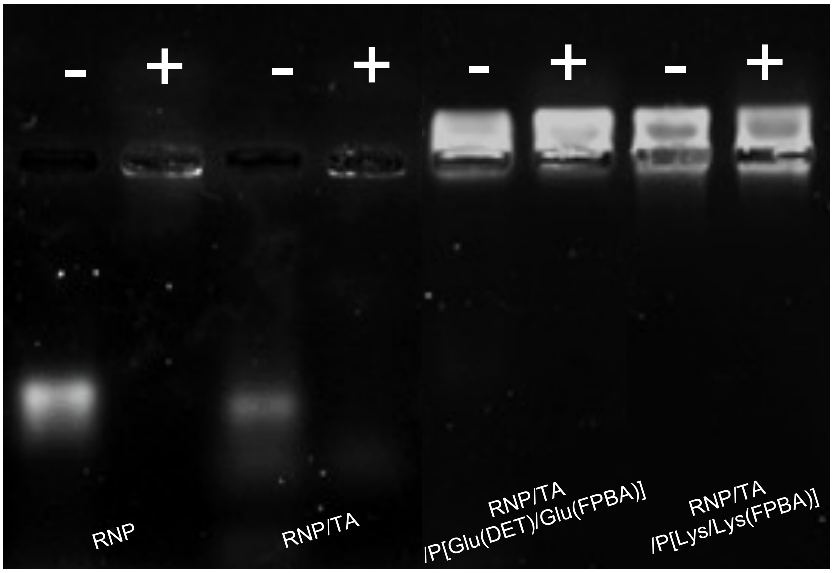


**Fig. S11.** RNase degradation stability detected by 1 % agarose gel electrophoresis. RNP concentration: 1 μM, [TA]/[RNP] = 30, [PBA-conjugated polymer]/[RNP] = 60. - and + means whether RNase treatment was done or not.

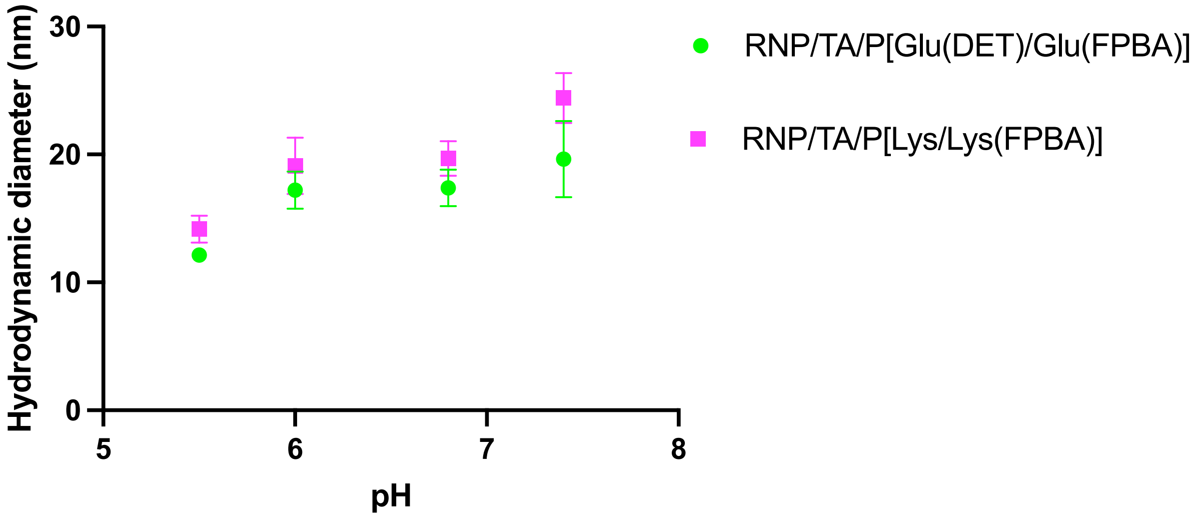


**Fig. S12.** pH-dependent hydrodynamic diameters of RNP ternary complexes. RNP concentrations: 200 nM, The molar ratio of [RNP]/[TA]/[PBA-conjugated polymer]= 1/30/30. The results are expressed as mean ±S.D. (n = 10).

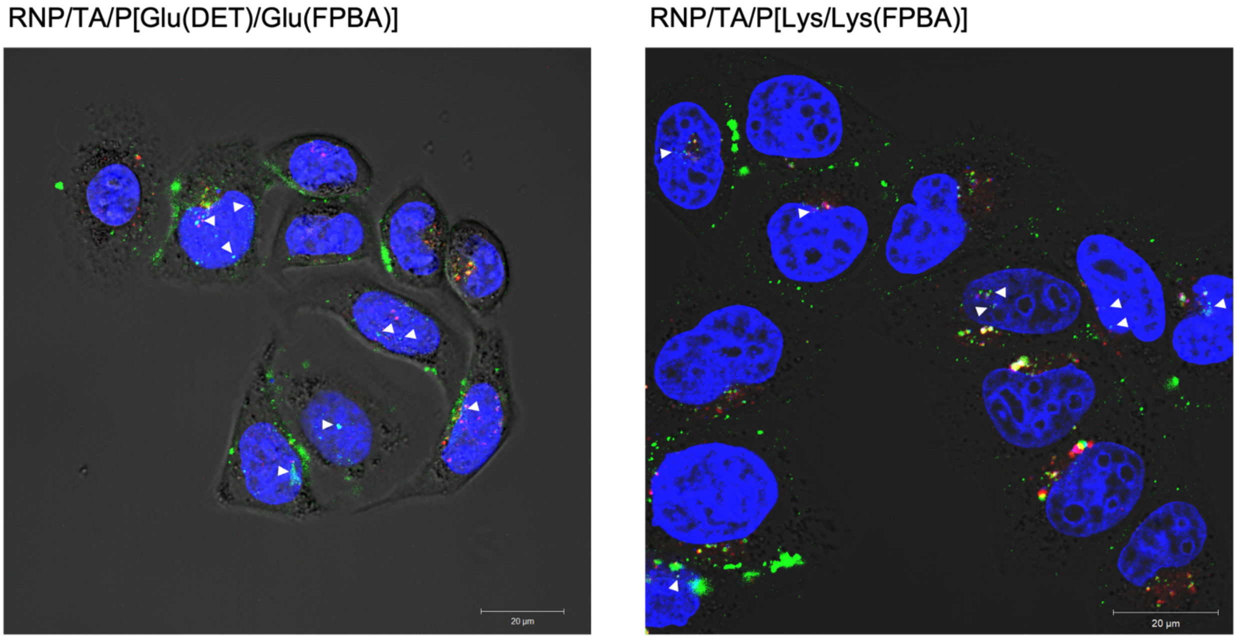


**Fig. S13.** Large scale subcellular distribution observed by CLSM. The Hela-Luc cells were incubated with the samples for 24 h. RNP, Lysotracker Red DND-99, and nucleus are shown in green, red, and blue respectively. Scale bar, 20 μm.

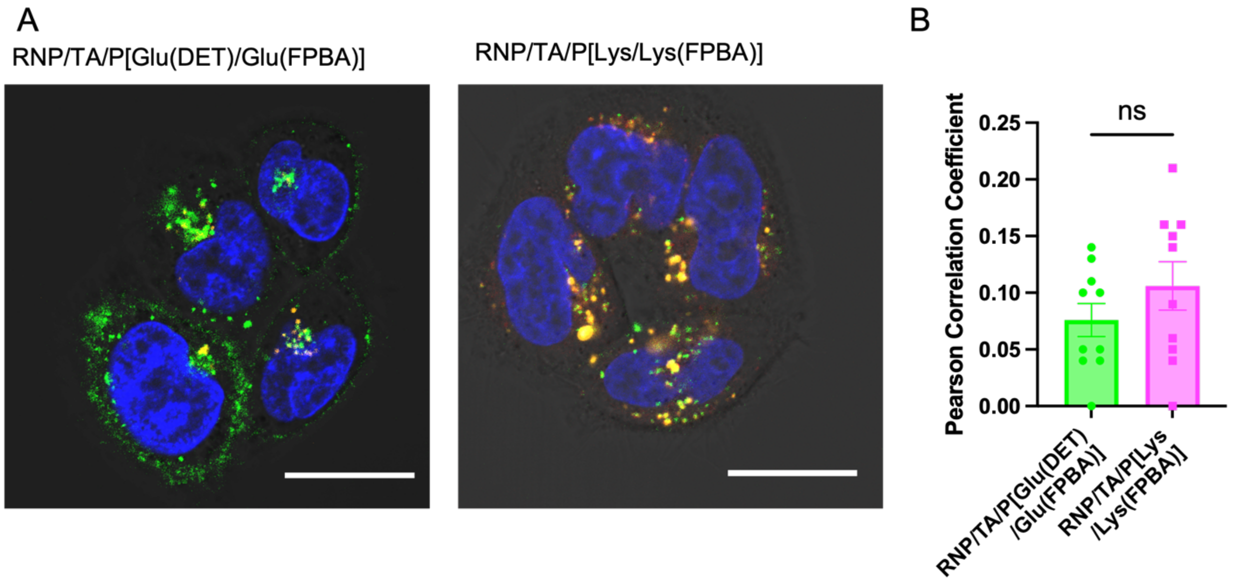


**Fig. S14.** CLSM image for confirmation of intracellular dissociation of the RNP ternary complex. The Hela-Luc cells were incubated with RNP ternary complexes containing Cas9-GFP and Alexa647-labelled polymers for 24 h. Cas9-GFP, Alexa647, and nucleus are shown in green, orange, and blue respectively. Scale bar, 20 µm. (B) Pearson's correlation coefficient of RNP with polymers. The results are expressed as mean ± S.E.M. (n = 10). Unpaired t-test was used for statistical analysis.

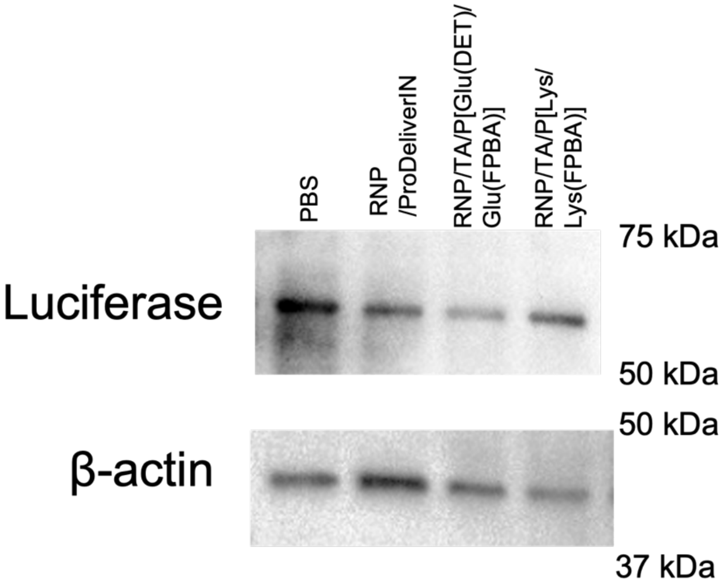


**Fig. S15.** Representative Western blot image showing Luc expression in Hela-Luc cells treated with PBS, RNP/ProDeliverIN, RNP/TA/P[Glu(DET)/Glu(FPBA)] and RNP/TA/P[Lys/Lys(FPBA)].

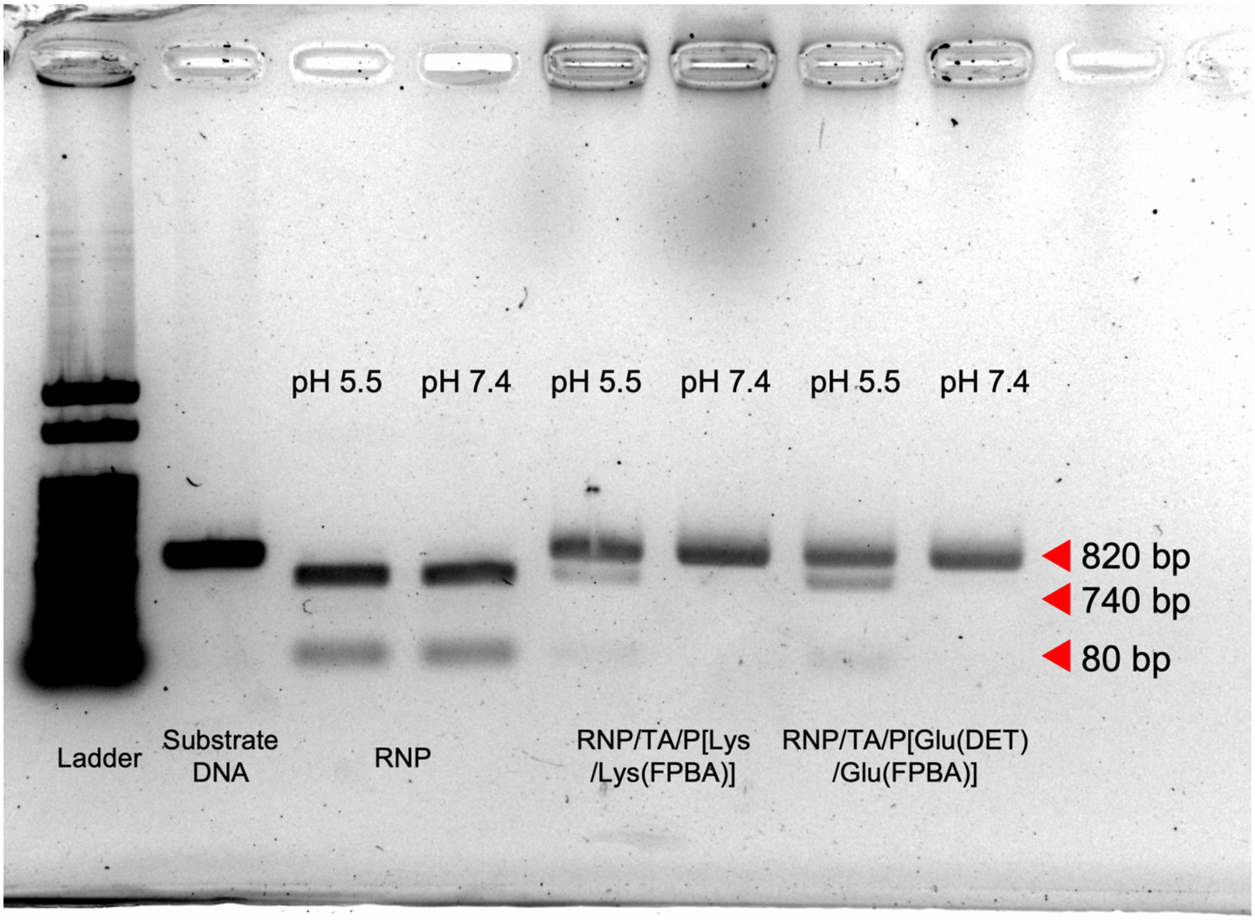


**Fig. S16.** *In vitro* DNA cleavage assay. The extracted Luc target region DNA with 300 ng was incubated with RNP samples in D-PBS(-) pH 7.4- or 50-mM MES buffer pH 5.5 for 60 min at 37°C. After the incubation, the mixture was neutralized by 50mM Tris buffer (pH 8.0) and DNA segments was isolated by agarose electrophoresis followed by SYBR safe staining for visualization. RNP concentration: 30 pmol, [RNP]/[TA]/[PBA-conjugated polymer] = 1/50/250.

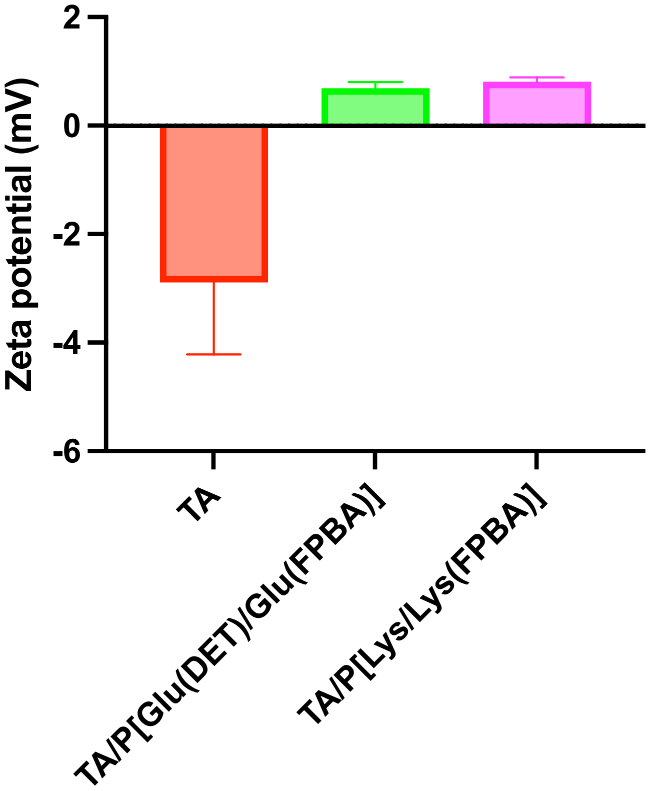


**Fig. S17.** Zeta potential of TA samples. The results are expressed as mean ±S.D. (n = 3).

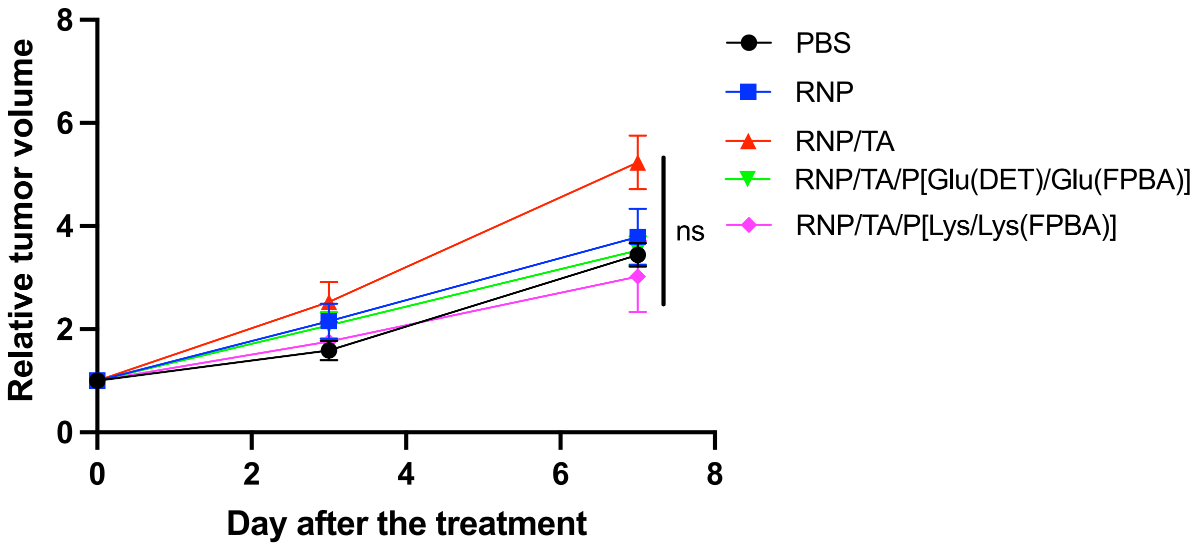


**Fig. S18.** Tumor size of subcutaneous Hela-Luc after RNP sample containing Luc-target sgRNA -treatment. The results are expressed as mean ±S.E.M. (n = 5).

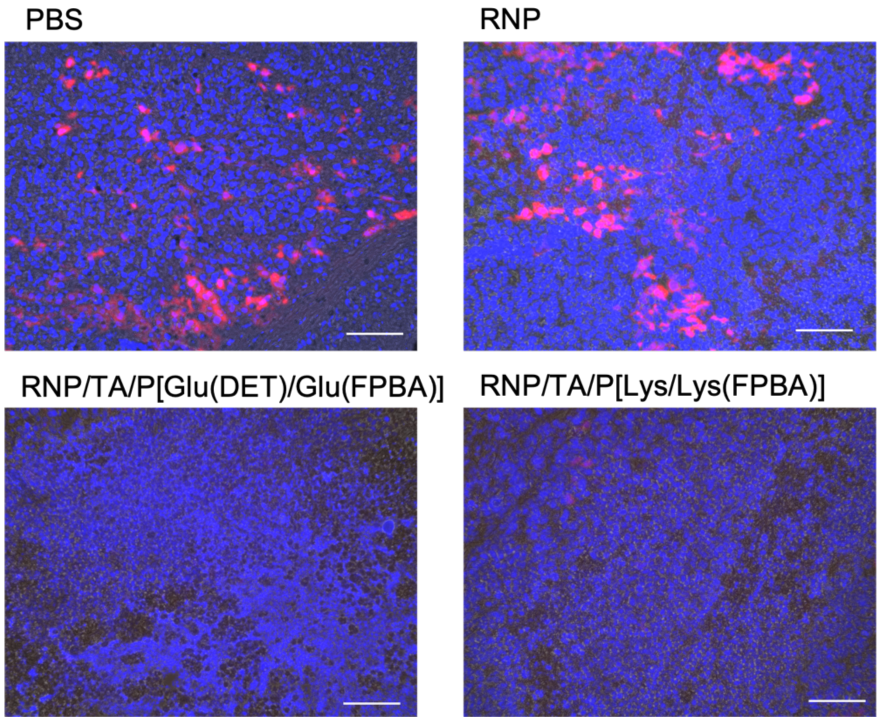


**Fig. S19.** Immunofluorescence image Hela-Luc tumors treated with PBS, RNP, RNP/TA/P[Glu(DET)/Glu(FPBA)] and RNP/TA/P[Lys/Lys(FPBA)]. Blue, Hoechst; Red, Luc. Scale bar, 100 μm.

A

B

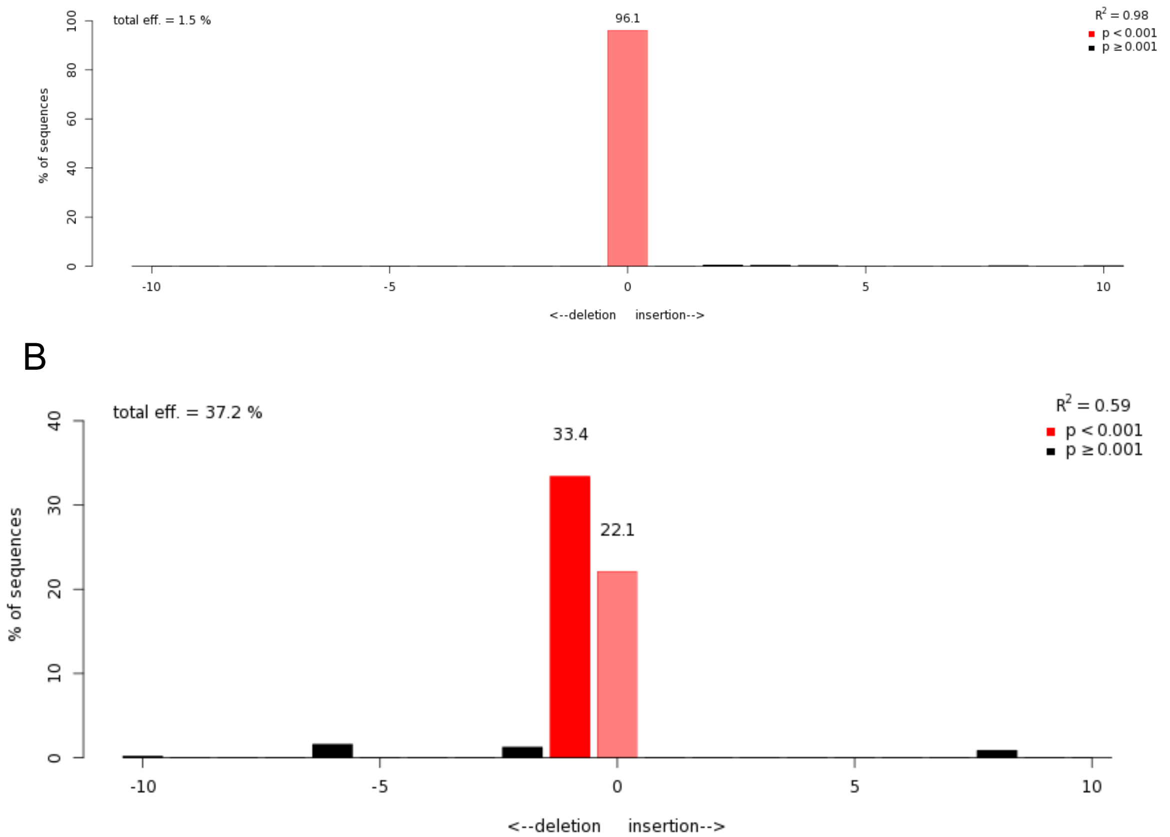


**Fig. S20.** TIDE analysis of sgLuc target loci of (A) RNP and (B) RNP/TA/P[Lys/Lys(FPBA)] treated group in *in vivo* Luc gene editing.

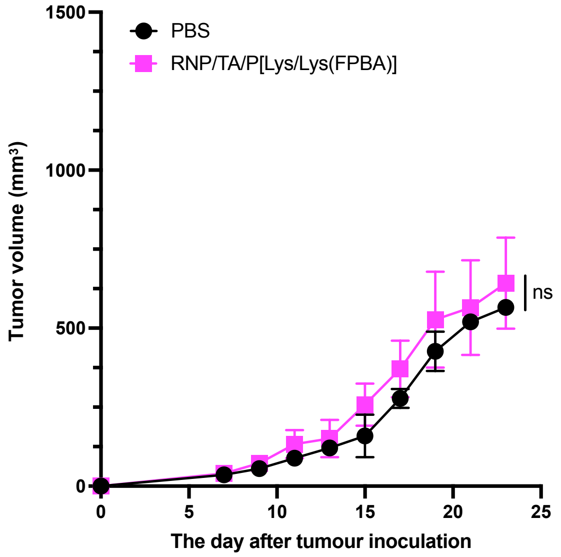


**Fig. S21.** The tumor growth curve of Hela-Luc tumor model. We inoculated Hela-Luc tumor cells subcutaneously on day 0. PBS and RNP/TA/P[Lys/Lys(FPBA)] were administrated on day 11, day 13, and day 15. RNP was formed of Cas9 and sgRNA targeting Luc. The result is expressed as mean ± S.E.M. (n = 5).

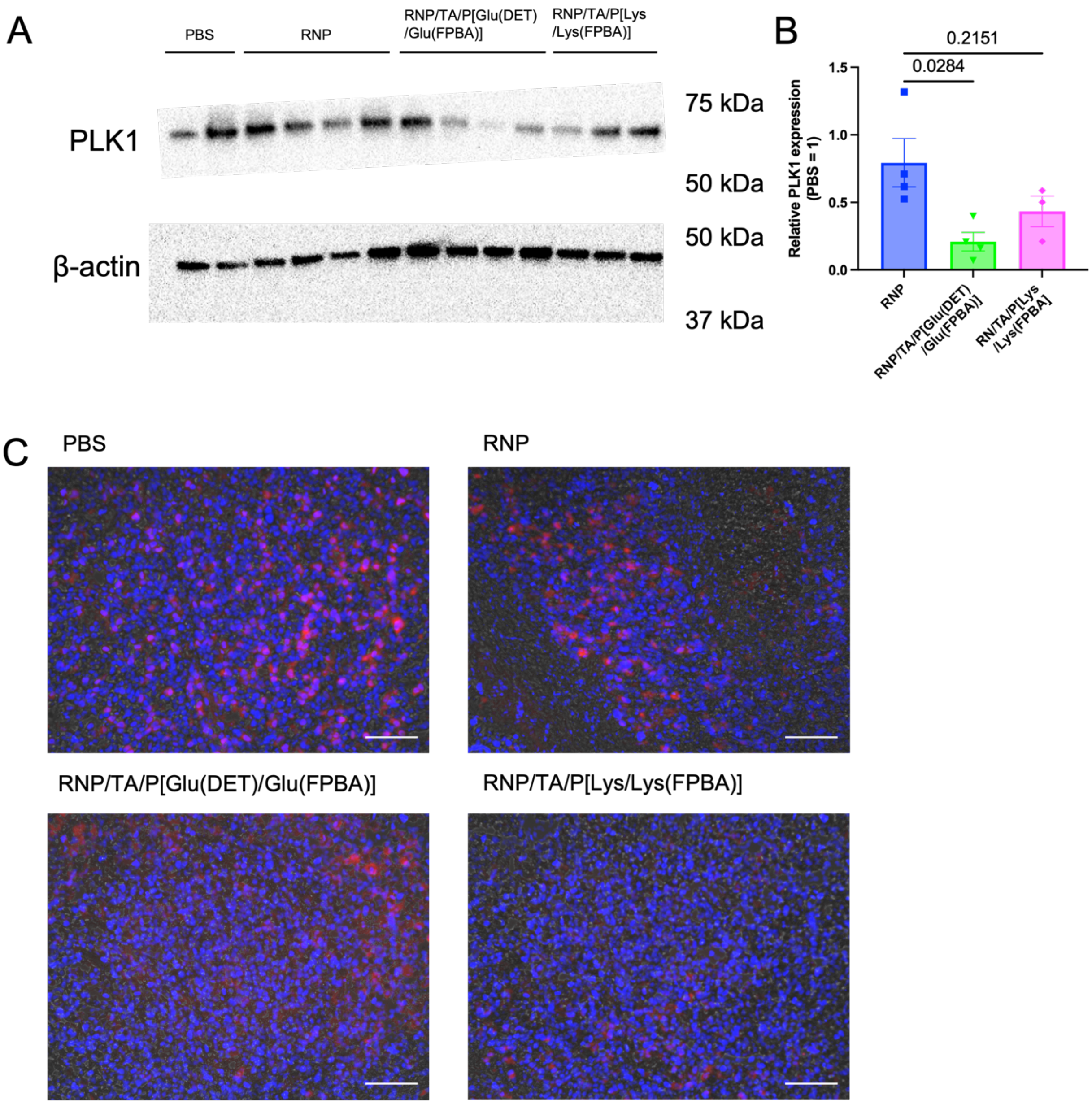


**Fig. S22.** *In vivo* PLK1 expression level in Hela-Luc tumors (A) Representative Western blot image showing PLK1 expression in Hela-Luc tumors treated with PBS and RNP samples (B) Densitometric quantification of Luc levels normalized to ß-Actin levels, and the relative values were then plotted against the PBS-treated group. Data are presented as mean ± SEM (n=3). Statistical significance was determined by ANOVA with Tukey's multiple comparison test (*p < 0.05). (C) Immunofluorescence image of Hela-Luc tumors treated with PBS, RNP, RNP/TA/P[Glu(DET)/Glu(FPBA)] and RNP/TA/P[Lys/Lys(FPBA)]. Blue, Hoechst; Red, PLK1. Scale bar, 100 μm.

A

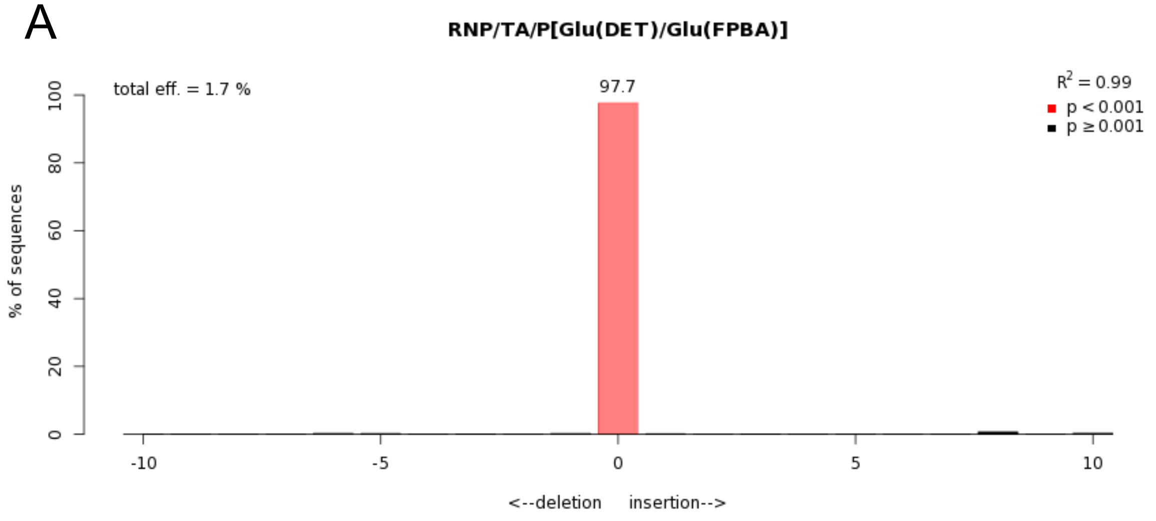


B

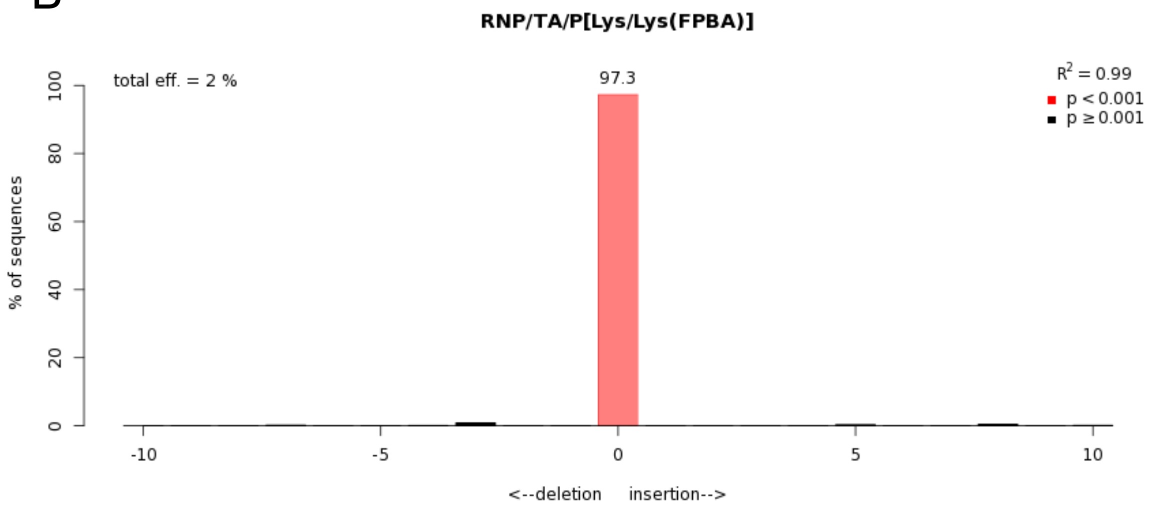


**Fig. S23.** TIDE analysis of sgPLK1 target loci of (A) RNP/TA/P[Glu(DET)/Glu(FPBA)] and (B) RNP/TA/P[Lys/Lys(FPBA)] treated group in *in vivo* PLK1 gene editing.

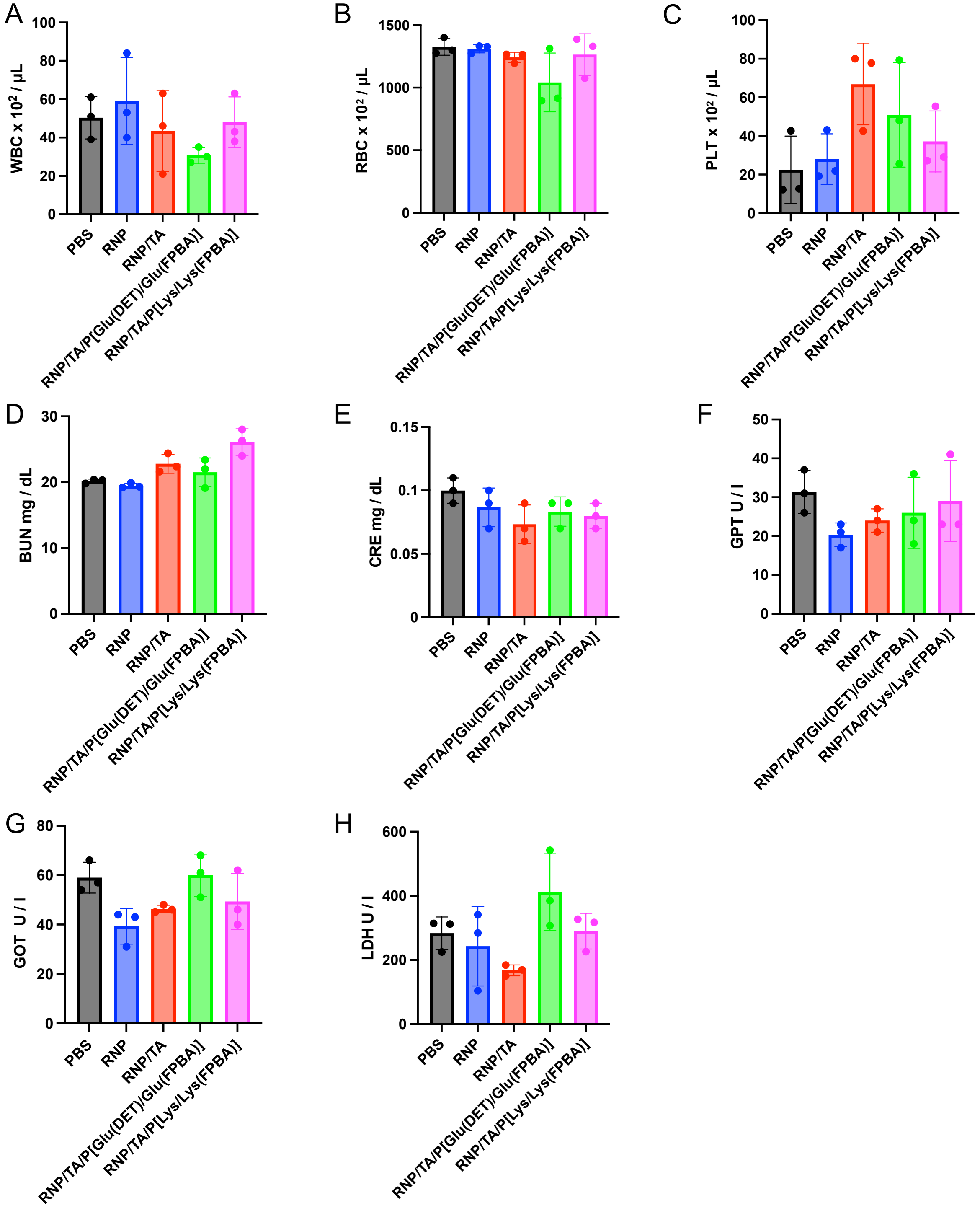


**Fig. S24.** Biochemical parameters of blood. (A) WBC, (B) RBC, (C) PLT, (D) BUN, (E) CRE, (F) GPT, (G) GOT/AST, and (H) LDH. Blood was collected 24h after intravenous injection of samples (Cas9; 32 µg/mouse, sgRNA; 6.4 µg/mouse, [RNP]/[TA]/[PBA-conjugated polymer] = 1/30/600. The results are expressed as means ± S.D. (n = 3). WBC; white blood cells, RBC; red blood cells, PLT; platelet, BUN; blood urea nitrogen, CRE; creatinine, GPT-P; glutamic pyruvic transaminase, GOT/AST; Glutamic Oxaloacetic Transaminase/Aspartate transaminase, LDH; lactic acid dehydrogenase.

**Table S1.**
$M_w/M_n$ of PEG-P[Lys(TFA)] and PEG-PBLG.

| Polymers | $M_W$ of PEG | DP | $M_w/M_n$ |
|---|---|---|---|
| PEG-P[Lys(TFA)] | 10,000 | 20 | 1.16 |
| PEG-PBLG | 10,000 | 24 | 1.08 |

**Table S2.**
Sequences of sgRNA used in this study.

| Nucleic acids | Target Sequences (5' to 3') | PAM (5' to 3') |
|---|---|---|
| Luciferase | CTTCGAAATGTCCGTTCGGT | TGG |
| PLK1 | CGGAGGCTCTGCTCGGATCG | AGG |
| KRAS$^{G13D}$ | GTTGGAGCTGGTGACGT | AGG |

**Table S3.**
Sequences of primers used in this study.

| Nucleic acids | Sequence (5' to 3') |
|---|---|
| Luciferase_F_vitro | AGAGCAACTGCATAAGGCTATGAAGAGATAC |
| Luciferase_R_vitro | CTTTAGGCAGACCAGTAGATCCAGAGGAGT |
| Luciferase_F_vivo | GGAACAATTGCTTTTACAGATGCACATATC |
| Luciferase_R_vivo | CGTGTAAATTAGATAAATCGTATTTGTCAATCA |
| PLK1_F_vivo | CGTGTCAATCAGGTTTTCCC |
| PLK1_R_vivo | TTGAGAAGCAGAGACTTAGG |
